# Daxx mediated histone H3.3 deposition on HSV-1 DNA restricts genome decompaction and the progression of immediate-early transcription

**DOI:** 10.1101/2024.08.15.608064

**Authors:** Ashley P.E. Roberts, Anne Orr, Victor Iliev, Lauren Orr, Steven McFarlane, Zhousiyu Yang, Ilaria Epifano, Colin Loney, Milagros Collados Rodriguez, Anna R. Cliffe, Kristen L. Conn, Chris Boutell

**Affiliations:** MRC-University of Glasgow Centre for Virus Research (CVR), Sir Michael Stoker Building, Garscube Campus, Glasgow, Scotland, UK; School of Life and Environmental Sciences, College of Health and Science, Joseph Banks laboratories, University of Lincoln, Brayford Pool Campus, Lincoln, LN6 7TS, UK; Department of Microbiology, Immunology and Cancer Biology, University of Virginia, Charlottesville, VA, USA; Department of Veterinary Microbiology, Western College of Veterinary Medicine, University of Saskatchewan, Saskatoon, SK, CAN

**Keywords:** HSV-1, histone, nucleosome, ICP0, Daxx, PML-NB

## Abstract

Herpesviruses are ubiquitous pathogens that cause a wide range of disease. Upon nuclear entry, their genomes associate with histones and chromatin modifying enzymes that regulate the progression of viral transcription and outcome of infection. While the composition and modification of viral chromatin has been extensively studied on bulk populations of infected cells by chromatin immunoprecipitation, this key regulatory process remains poorly defined at single-genome resolution. Here we use high-resolution quantitative imaging to investigate the spatial proximity of canonical and variant histones at individual Herpes Simplex Virus 1 (HSV-1) genomes within the first 90 minutes of infection. We identify significant population heterogeneity in the stable enrichment and spatial proximity of canonical histones (H2A, H2B, H3.1) at viral DNA (vDNA) relative to established promyelocytic leukaemia nuclear body (PML-NB) host factors that are actively recruited to viral genomes upon nuclear entry. We show the replication-independent histone H3.3/H4 chaperone Daxx to cooperate with PML to mediate the enrichment and spatial localization of variant histone H3.3 at vDNA that limits the rate of HSV-1 genome decompaction to restrict the progress of immediate-early (IE) transcription. This host response is counteracted by the viral ubiquitin ligase ICP0, which degrades PML to disperse Daxx and variant histone H3.3 from vDNA to stimulate the progression of viral genome expansion, IE transcription, and onset of HSV-1 replication. Our data support a model of intermediate and sequential histone assembly initiated by Daxx that limits the rate of HSV-1 genome decompaction independently of the stable enrichment of histones H2A and H2B at vDNA required to facilitate canonical nucleosome assembly. We identify HSV-1 genome decompaction upon nuclear infection to play a key role in the initiation and functional outcome of HSV-1 lytic infection, findings pertinent to the transcriptional regulation of many nuclear replicating herpesvirus pathogens.

## Introduction

Herpesviruses are a ubiquitous family of pathogens that cause a variety of clinically important diseases, ranging from mild skin sores and rashes to severe birth defects, cancer, and life-threatening encephalitis [1]. A common feature shared by all herpesviruses is the configuration of their double-stranded DNA genomes that range in size from 125 to 240 kb in length [2]. These genomes are tightly packaged into viral capsids devoid of cellular protein (e.g., histones) under extreme pressure in the presence of spermine [3–11]. Following entry, these genomes are delivered into the nucleus of newly infected cells [12–15], where they appear as compact foci (∼ 0.1 μm^3^) that progressively expand following the initiation of transcription into viral DNA (vDNA) replication centres [16, 17].

Compaction of eukaryotic chromatin occurs through nucleosome formation, with each nucleosome comprising of an octamer of histones (two molecules each of H2A, H2B, H3.1, and H4) wrapped in approximately 147 bp of DNA. Nucleosome assembly can occur in a DNA replication-dependent and -independent manner through the specific loading functions of histone chaperones [18–21]. The expression of canonical histones is robustly induced during S-phase. In contrast, histone variants (including histone H3.3, H2A.Z, macroH2A, and H2A.X) are constitutively expressed throughout the cell cycle and actively exchanged into chromatin to define specific chromatin boundaries [22]. The organization of cellular chromatin into euchromatin (transcriptionally active) or heterochromatin (transcriptionally repressive) is controlled by histone reader complexes that bind specific histones carrying distinct epigenetic modifications that regulate gene accessibility and DNA compaction state [21]. While chromatin immunoprecipitation (ChIP) studies have demonstrated Herpes Simplex Virus 1 (HSV-1) genomes to bind histones carrying epigenetic marks indicative of both euchromatin (H3K4me3, acetylated H3) and heterochromatin (H3K9me2/me3 and H3K27me2/me3) dependent on the presence or absence of viral transactivating proteins (e.g., VP16 or ICP0) during productive infection [23–27], the *de novo* assembly of chromatin on viral DNA (vDNA) remains poorly defined on a genome population basis. Consequently, it remains unclear as to whether the composition or epigenetic modification of viral chromatin identified by ChIP is representative of the total population of genomes under investigation or a limited subset of enriched genomes. Thus, it remains to be determined to what degree population heterogeneity in viral chromatin assembly or epigenetic modification may functionally contribute to the outcome of infection.

We and others have shown HSV-1 genomes pre-labelled with EdC (5-Ethynyl-2’-deoxycytidine) to enable the single-molecule detection of vDNA by click chemistry [16, 17, 27–29]. We have shown nuclear infecting HSV-1 genomes to rapidly associate with core constituent proteins of promyelocytic leukaemia nuclear bodies (PML-NBs) leading to vDNA entrapment [16]. This nuclear host defence to infection is counteracted by the HSV-1 ubiquitin ligase ICP0 [30–32], which induces the proteasome-dependent degradation of PML (the major scaffolding protein of PML-NBs, [33]), leading to the dispersal of repressive PML-NB host factors from vDNA that stimulates the onset of lytic infection [31, 32, 34–37].

HSV-1 nuclear infection also induces the recruitment of two DNA histone H3.3/H4 chaperones to vDNA; Daxx (death domain associated protein 6) and HIRA (histone cell cycle regulator) [16, 38–41]. The stable recruitment of these histone chaperones to vDNA occurs asynchronously, with Daxx localizing to genomes immediately upon nuclear entry [16, 39], while HIRA recruitment is dependent on the initiation of vDNA replication or stimulation of cytokine-mediated immune defences [38, 42, 43]. Notably, both Daxx and HIRA colocalize with PML at vDNA, which can lead to the epigenetic modification of histone H3.3 indicative of heterochromatin silencing [27, 28, 42–44]. Recent evidence also has shown PML to influence the equilibrium of heterochromatic marks associated with vDNA (e.g., H3K27me2 *vs*. H3K9me3) [27] and viral genomes associated with PML-NBs to be more transcriptionally repressed during viral latency [44]. Such observations have led to the hypothesis that PML-NBs may act as sites for the assembly and/or maintenance of viral heterochromatin.

However, it remains to be formally investigated as to what role PML-NB entrapment of vDNA may play in the *de novo* assembly of viral chromatin upon nuclear infection. We therefore set out to investigate the spatial localization and relationship between PML-NB entrapment of HSV-1 genomes and vDNA chromatin assembly using quantitative imaging at single-genome resolution.

We identify significant population heterogeneity in the stable enrichment and spatial proximity of canonical histones (H2A, H2B, and H3.1) to nuclear infecting HSV-1 genomes entrapped within PML-NBs. We show PML-NBs not to sterically inhibit the enrichment of these canonical histones to infecting genomes, but to cooperate with Daxx in the spatial localization and enrichment of variant histone H3.3 at vDNA. We show HSV-1 genomes released from capsids *in vitro* to have equivalent volumetric dimensions to those observed inside infected cells upon nuclear infection, demonstrating HSV-1 genomes to retain a significant degree of vDNA compaction post-capsid release independently of chromatin assembly. We demonstrate Daxx to be responsible for the localization of variant histone H3.3 at vDNA that limits the rate of HSV-1 genome decompaction and progression of viral immediate-early (IE) transcription. This host response to nuclear infection is counteracted by the HSV-1 ubiquitin ligase ICP0, which degrades PML to disperse Daxx and variant histone H3.3 from vDNA to stimulate genome expansion, progression of IE transcription, and the onset of viral lytic replication. Our study identifies Daxx as a key mediator in the intermediate and sequential assembly of viral chromatin that constrains HSV-1 genome decompaction post-capsid release independently of stable canonical nucleosome assembly.

## Results

### Canonical histones do not stoichiometrically localize to nuclear infecting HSV-1 genomes

We began our analysis by validating HSV-1 genomes to bind cellular histones upon nuclear infection. Human foreskin fibroblast (HFt) cells were infected with an HSV-1 ICP0-null mutant (ΔICP0; MOI 3 PFU/cell) to prevent the ICP0-dependent disruption of viral chromatin and proteasome-dependent degradation of PML-NBs [30, 45]. Cells were harvested at 90 minutes post-infection (90 mpi; post-addition of virus) and viral chromatin immunoprecipitated (IP) using ChIP-grade histone antibodies or species-matched IgG (negative control). Consistent with previous studies [45–47], IP of histones led to the recovery of HSV-1 DNA (Fig. 1A), demonstrating a proportion (∼ 0.5 to 1 % of soluble input) of infecting genomes to stably bind histones. We hypothesized that if nucleosome assembly were required to promote HSV-1 genome compaction post-capsid release that we would observe the stable enrichment of canonical histones (H2A, H2B, H3.1, and H4) at vDNA on a population wide basis [20, 48]. As PML-NBs rapidly entrap infecting HSV-1 genomes [16], and are known repositories for variant histone H3.3 deposition [49–51], we first examined the localization of histones at PML-NBs in mock-treated cells. For controls, we examined the localization of two histone H3.3/H4 chaperones, Daxx and HIRA, known to either reside or transiently associate with PML-NBs, respectively [33, 38, 42, 52, 53]. We observed Daxx, but not HIRA, to stably localize at PML-NBs in mock-treated cells (Fig. 1B, S1). Little to no stable colocalization of canonical histones H2A or H2B were observed at PML-NBs (Fig. 1B), which predominantly localized to cellular chromatin interspersed with bright puncta (Fig. S1). Using an antibody that recognized both canonical (H3.1) and variant (H3.3) histone H3 (Fig. S2), we observed consistent histone H3 localization at PML-NBs (Fig. 1B, S1).

**Fig 1.**
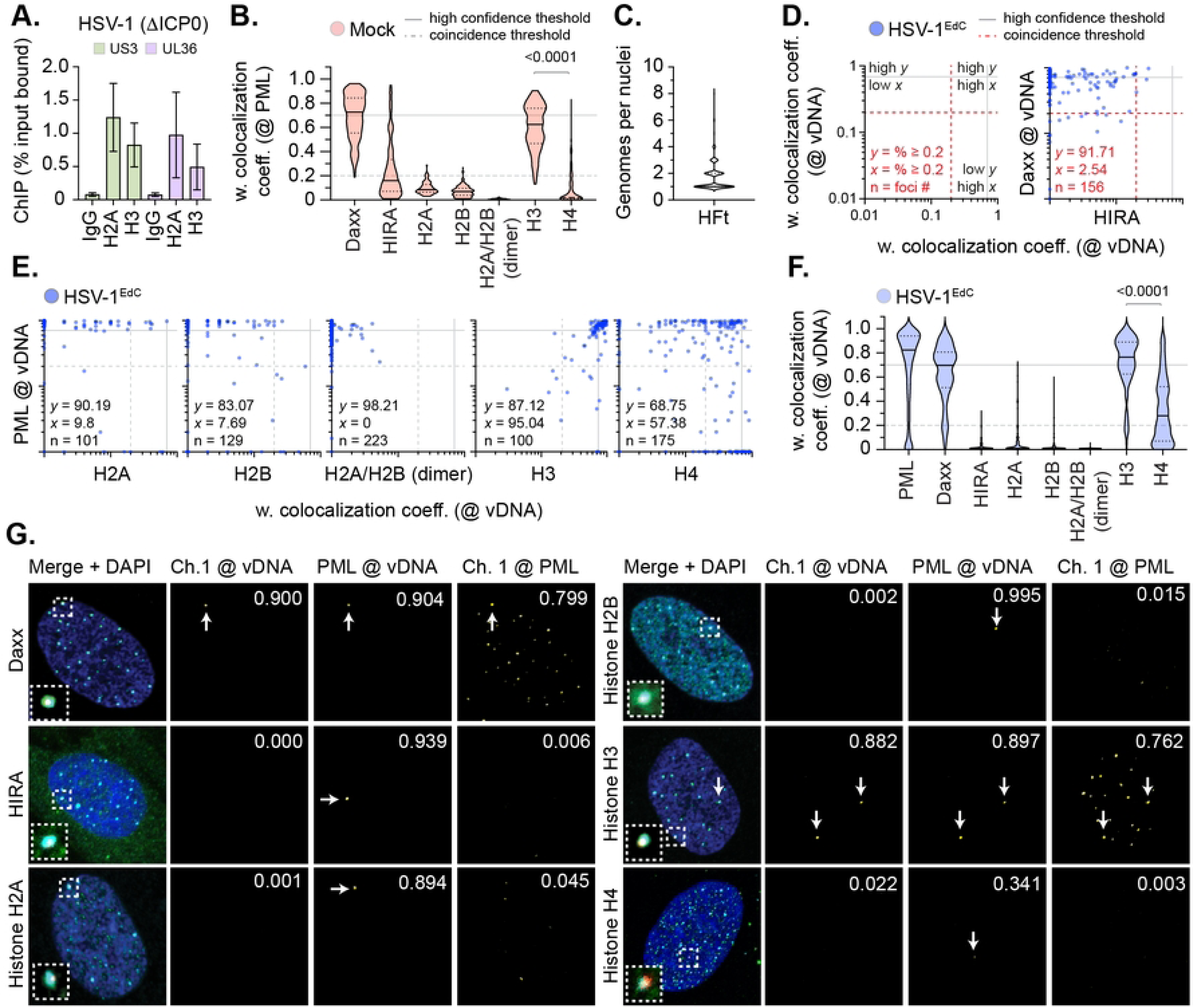
Canonical histones do not stoichiometrically localize to nuclear infecting HSV-1 genomes. (A) HFt cells were infected with an HSV-1 ICP0-null mutant (ΔICP0; MOI of 3 PFU/cell). Chromatin extracts were prepared at 90 mins post-infection (mpi; post-addition of virus) and subjected to ChIP using ChIP-grade anti-histone H2A or histone H3 antibodies and species-matched IgG (negative control). Bound viral DNA (vDNA) was quantified by qPCR using probes specific to HSV-1 US3 or UL36. Values were normalized to input loading controls and presented as percentage (%) input bound. Means and SEM shown. (B to G) HFt cells were mock-treated or infected with WT HSV-1^EdC^ (MOI of 1 PFU/cell). Cells were fixed at 90 mpi and stained for PML, Daxx, HIRA, histones H2A, H2B, H2A/H2B heterodimers (dimer), H3, or H4 by indirect immunofluorescence. vDNA was detected by click chemistry [16]. Nuclei were stained with DAPI. (B) Colocalization frequency of cellular proteins at PML-NBs in mock-treated HFt cells. N≥60 nuclei per staining condition. Confocal microscopy images shown in Fig. S1. Violin plots: median weighted (w.) colocalization coefficient (coeff.), solid black line; 25^th^ to 75^th^ percentile range, dotted black lines; coincidence threshold (0.2), dotted grey line; high confidence threshold (0.7), solid grey line. Threshold values determined from scatter plots shown in Fig. 1D (Daxx positive control/HIRA negative control) and Fig. 2B (PML positive control/eYFPnls negative control) [16, 38]. Mann-Whitney *U*-test, *P*-value shown. (C) Quantitation of the number of genome foci detected in the nucleus of HSV-1^EdC^ infected cells at 90 mpi. N=883 nuclei derived from 18 independent experiments. (D/E) Scatter plots showing paired w. colocalization coeff. values of proteins of interest (indicated on *x*- and *y*-axis) at vDNA. Percentage (%) of genomes ≥ coincident threshold (*x/y* ≥ 0.2) per sample condition shown; number (n) of genome foci analysed per sample population shown. (F) Colocalization frequency of cellular proteins of interest at vDNA in HSV-1 infected HFt cells at 90 mpi (as in D, E). Mann-Whitney *U*-test, *P*-value shown. (G) Merged images of Daxx, HIRA, histones H2A, H2B, H3, or H4 (Channel 1 (Ch.1), green; as indicated), and PML (cyan) colocalization at vDNA (red). Cut mask (yellow) highlights regions of colocalization between cellular proteins of interest and vDNA or PML (as indicated); w. colocalization coeff. shown. Dashed boxes show magnified regions of interest. White arrows highlight regions of colocalization at vDNA. Individual panels shown in Fig. S4. (A to G) Data derived from a minimum of three independent experiments. Raw values presented in S1 data.

Analysis of transgenic HFt cell lines induced to express fluorescently tagged (mEmerald; mEm) histones corroborated this colocalization to be specific for variant histone H3.3 (Fig. S2, H3.3-mEm; [49–51]). No colocalization was observed for histones H2A-mEm, H2B-mEm, and H3.1-mEm at PML-NBs (Fig. S2). In contrast to the detection of endogenous histone H4 (Fig. 1B, S1), ectopic expression of histone H4-mEm led to its consistent detection at PML-NBs to levels equivalent to that of H3.3-mEm (Fig. S2). We posit that the endogenous detection of histone H4 at PML-NBs may be subject to epitope masking when in complex with Daxx at PML-NBs [54]. Importantly, all mEm-tagged histones could be observed to associate with mitotic cellular chromatin (Fig. S3), demonstrating that the fusion of the mEm tag onto the C-terminus of each histone not to impair nucleosome assembly. We conclude canonical histones H2A, H2B, and H3.1 not to be stably enriched at PML-NBs prior to HSV-1 infection.

We next examined the localization of endogenous histones to nuclear infecting HSV-1 genomes by click chemistry and indirect immunofluorescence. Cell monolayers were infected with WT HSV-1^EdC^ (MOI of 1 PFU/cell) and fixed at 90 mpi, a time point in the linear phase of genome delivery to the nucleus (median frequency of 1 genome per nucleus, Fig 1C) [16]. As expected, PML and Daxx robustly localized at vDNA, while HIRA did not (Fig. 1D to G, S4) [16, 38]. The frequency of histone H2A and H2B colocalization at vDNA was equivalent to that of HIRA (Fig. 1E, F, S4), with few genomes showing colocalization at 90 mpi. Immunolabelling using a fluorescent H2A/H2B nanobody, which readily detected H2A/H2B heterodimers associated with cellular chromatin (Fig. S5), also failed to detect these histones to be stably enriched at vDNA (Fig. 1E, F, S5; H2A/H2B dimer). In contrast, histone H3 (H3.1/H3.3) colocalized at vDNA with a frequency equivalent to that of Daxx (Fig. 1D to G, S4). While histone H4 was observed to be enriched at vDNA, the frequency of this colocalization was lower than that observed for histone H3 (Fig. 1E, F). Analysis of paired data demonstrated a substantial degree of population heterogeneity in the colocalization of histones at vDNA relative to PML (positive control; Fig. 1E), with histones H3 and H4 exhibiting the highest frequencies of association on a genome population basis at 90 mpi (Fig. 1E, F).

To exclude the possibility of differences in antibody avidity influencing the detection of histones at vDNA, we next examined the localization of mEm-tagged histones at vDNA under equivalent infection conditions. Only histones H3.3-mEm and H4-mEm stably localized at vDNA, with little to no colocalization observed for eYFPnls (negative control) or histones H2A-mEm, H2B-mEm, or H3.1-mEm (Fig. 2A to C, S6). Analysis of paired data again demonstrated a substantial degree of population heterogeneity in the colocalization of canonical H2A-mEm, H2B-mEm, and H3.1-mEm histones at vDNA relative to PML (Fig. 2B). Thus, a failure to detect the stable enrichment of these histones at vDNA is not a consequence of epitope masking or relative differences in antibody avidity. ChIP analysis of infected cells using an anti-GFP antibody led to a similar profile of vDNA recovery to that observed for endogenous histones (Fig. 1A *vs*. 2D; ∼ 1 to 4 % total soluble input). Together, these data indicate that only a minor fraction of input genomes to stably bind histones on a genome population basis. Thus, the relative high frequency of colocalization observed for variant histone H3.3 and histone H4 at vDNA might relate to their reported interaction with Daxx at PML-NBs independently of their *de novo* deposition on vDNA (Fig. S1, S2; [49–51]).

**Fig 2.**
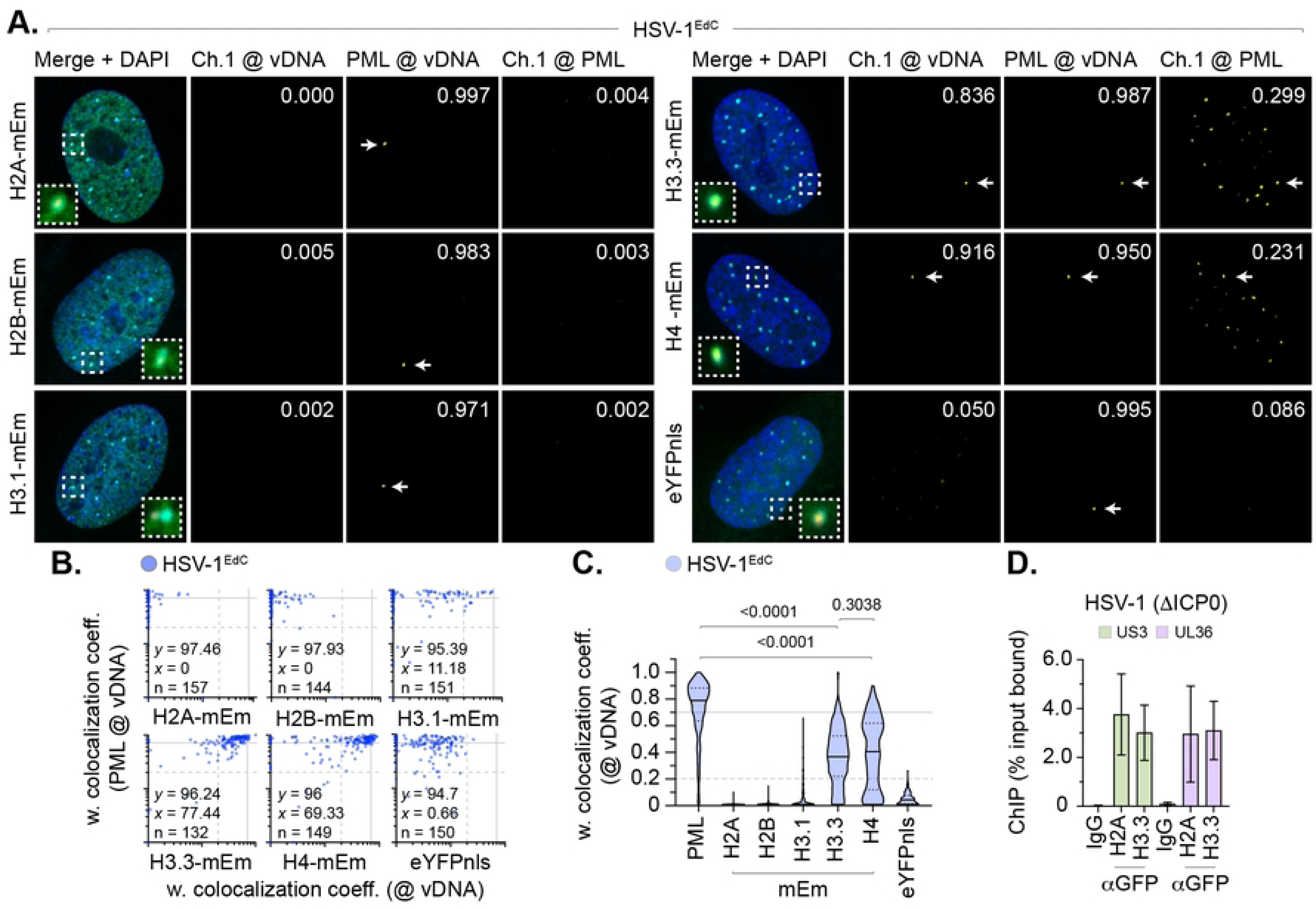
Fluorescent histones do not stoichiometrically localize to nuclear infecting HSV-1 genomes. (A to C) HFt cells were stably transduced with doxycycline inducible lentiviral vectors encoding C-terminally tagged fluorescent (mEmerald, mEm) histones or eYFPnls (negative control) as indicated. Cells were induced to express proteins of interest for 6 h prior to infection with WT HSV-1^EdC^ (MOI of 1 PFU/cell). Cells were fixed at 90 mpi and stained for PML by indirect immunofluorescence and vDNA by click chemistry. Nuclei were stained with DAPI. (A) Merged confocal microscopy images of mEm-tagged histones or eYFPnls (green) and endogenous PML (cyan) colocalization at vDNA (red). Cut mask (yellow) highlights regions of colocalization between cellular proteins of interest and vDNA or PML; weighted (w.) colocalization coefficient (coeff.) shown. Dashed boxes show magnified regions of interest. White arrows highlight regions of colocalization at vDNA. Individual panels shown in Fig. S6. (B) Scatter plots showing paired w. colocalization coeff. values of proteins of interest (indicated on *x*- and *y*-axis) at vDNA. Percentage (%) of genomes ≥ coincident threshold (*x/y* ≥ 0.2) per sample condition shown; number (n) of genome foci analysed per sample condition shown. (C) Colocalization frequency of proteins of interest at vDNA (as in B). Violin plots: median w. colocalization coeff., solid black line; 25^th^ to 75^th^ percentile range, dotted black lines; coincidence threshold (0.2), dotted grey line; high confidence threshold (0.7), solid grey line. Mann-Whitney *U*-test, *P*-values shown. (D) HFt cells were infected with an HSV-1 ICP0-null mutant (ΔICP0; MOI of 3 PFU/cell). Chromatin extracts were prepared at 90 mpi and subjected to ChIP using anti-GFP antibody or species-matched IgG (negative control). Bound viral DNA (vDNA) was quantified by qPCR using probes specific to HSV-1 US3 or UL36. Values were normalized to input loading controls and presented as percentage (%) input bound. Means and SEM shown. (A to D) Data derived from a minimum of three independent experiments. Raw values presented in S1 data.

To investigate whether the colocalization of histone H3.3 at vDNA correlated with its constitutive localization at PML-NBs and/or *de novo* deposition on viral genomes, we screened a panel of cell lines for endogenous histone H3.3 colocalization at PML-NBs. We identified HaCaT (human skin keratinocyte) cells to have reduced levels of variant histone H3.3 at PML-NBs in mock-treated cells (Fig. 3A, B) compared to other cell lines (HFt, HEL, RPE). This was surprising, as Daxx (the histone chaperone responsible for histone H3.3/H4 deposition at PML-NBs; [49–51]) and its binding partner ATRX (α-thalassemia mental retardation X-linked protein) were both robustly detected at PML-NBs (Fig. 3A, C). We posit the lack of constitutive histone H3.3 localization at PML-NBs may relate to the transformed and/or aneuploid nature of HaCaT cells [55]. Infection of HaCaT cells led to the stable enrichment of PML, histone H3 (H3.1/H3.3), and H4 (to a lesser extent) at vDNA (Fig. 3D, E). Whereas histones H2A and H2B did not (Fig. 3D, E). These data identify histones H3.3 and H4 to be actively recruited to vDNA independently of their sub-cellular localization at PML-NBs prior to infection. Taken together with our HFt analysis (Fig. 1, 2), these data indicate nuclear infecting HSV-1 genomes to be preferentially enriched for histones H3.3 and H4 on a population wide basis independently of the stable enrichment of H2A or H2B.

**Fig 3.**
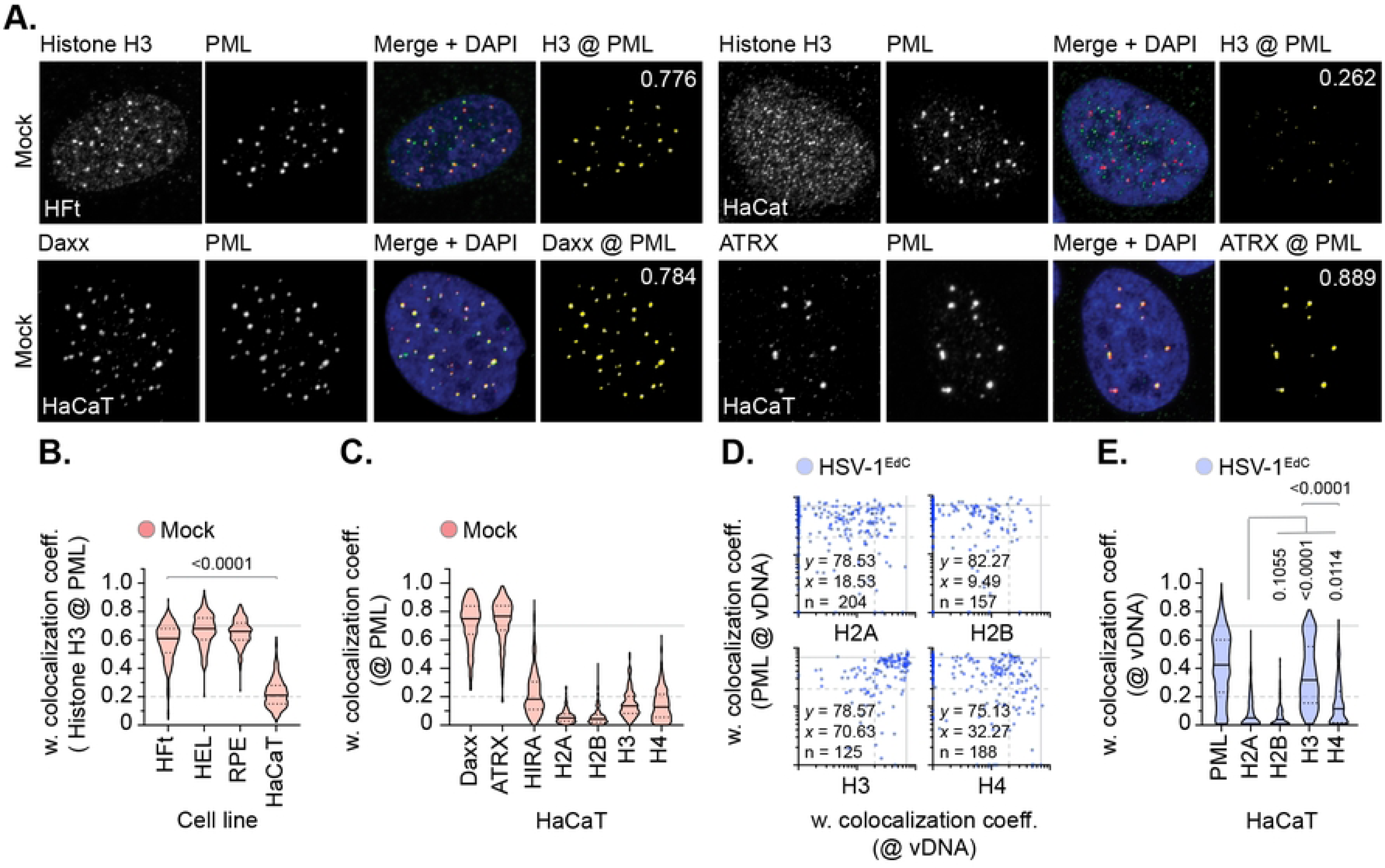
Histone H3 is enriched at vDNA independently of its sub-cellular localization at PML-NBs. (A to E) HFt, HEL, RPE, or HaCaT cells were mock-treated or infected with WT HSV-1^EdC^ (MOI of 1 PFU/cell). Cells were fixed at 90 mpi and stained for proteins of interest (as indicated) by indirect immunofluorescence and vDNA by click chemistry. Nuclei were stained with DAPI. (A) Confocal microscopy images of mock-treated HFt and HaCaT cells showing sub-nuclear localization of histone H3, Daxx, or ATRX (green) and PML (red). Cut mask (yellow) highlights regions of colocalization between cellular proteins of interest and PML; weighted (w.) colocalization coefficient (coeff.) shown. (B) Colocalization frequency of histone H3 at PML-NBs in mock-treated HFt, HEL, RPE, and HaCaT cells. Violin plots: median w. colocalization coeff., solid black line; 25^th^ to 75^th^ percentile range, dotted black lines; coincidence threshold (0.2), dotted grey line; high confidence threshold (0.7), solid grey line. Mann-Whitney *U*-test, *P*-value shown. N>240 nuclei per sample condition. (C) Colocalization frequency of proteins of interest at PML-NBs in mock-treated HaCaT cells. N≥250 nuclei per sample condition. (D) Scatter plots showing paired w. colocalization coeff. values of proteins of interest (indicated on *x*- and *y*-axis) at vDNA within infected HaCaT cells. Percentage (%) of genomes ≥ coincident threshold (*x/y* ≥ 0.2) per sample condition shown; number (n) of genome foci analysed per sample condition shown. (E) Distribution of PML, histones H2A, H2B, H3, and H4 colocalization frequency at vDNA (as in D). Mann-Whitney *U*-test (top), one-way ANOVA Kruskal-Wallis test (bottom), *P*-values shown. (A to E) Data derived from a minimum of three independent experiments. Raw values presented in S1 data.

### High-resolution imaging identifies canonical histones to localize in variable proximity to vDNA

We next investigated the spatial proximity of histones at vDNA utilizing high-resolution confocal microscopy imaging. All histones (H2A, H2B, H3.1/H3.3, and H4) could be observed to localize in relative proximity (2 μm^3^) to vDNA, which consistently demonstrated a high frequency of entrapment within PML-NBs (Fig. 4A) [16]. A subset of histones could be observed to make surface contact with PML and vDNA entrapped therein, with histone H3 (H3.1/H3.3) being in closest proximity to vDNA (Fig. 4B, C). Notably, histone H3 was observed to localize asymmetrically to both PML-NBs and vDNA entrapped therein (Fig. 4A, white arrows). Volumetric measurements demonstrated PML-NBs that contained vDNA to increase in size relative to non-genome containing PML-NBs within the same nucleus (Fig. 4D), indicative of an increase in PML-NB expansion upon vDNA entrapment. ChIP analysis identified PML to bind vDNA (Fig. 4E), identifying PML to be an accessory component of viral chromatin upon nuclear infection. Collectively, these data demonstrate canonical histones H2A and H2B to be proximal to vDNA on an individual genome basis, but to be spatially separate and distinct from histone H3.3 or PML on a genome population basis (Fig. 4A, C). Moreover, these data also suggest that a significant fraction of histone H3.3 localized at PML-NBs not to be directly associated with viral chromatin at 90 mpi.

**Fig 4.**
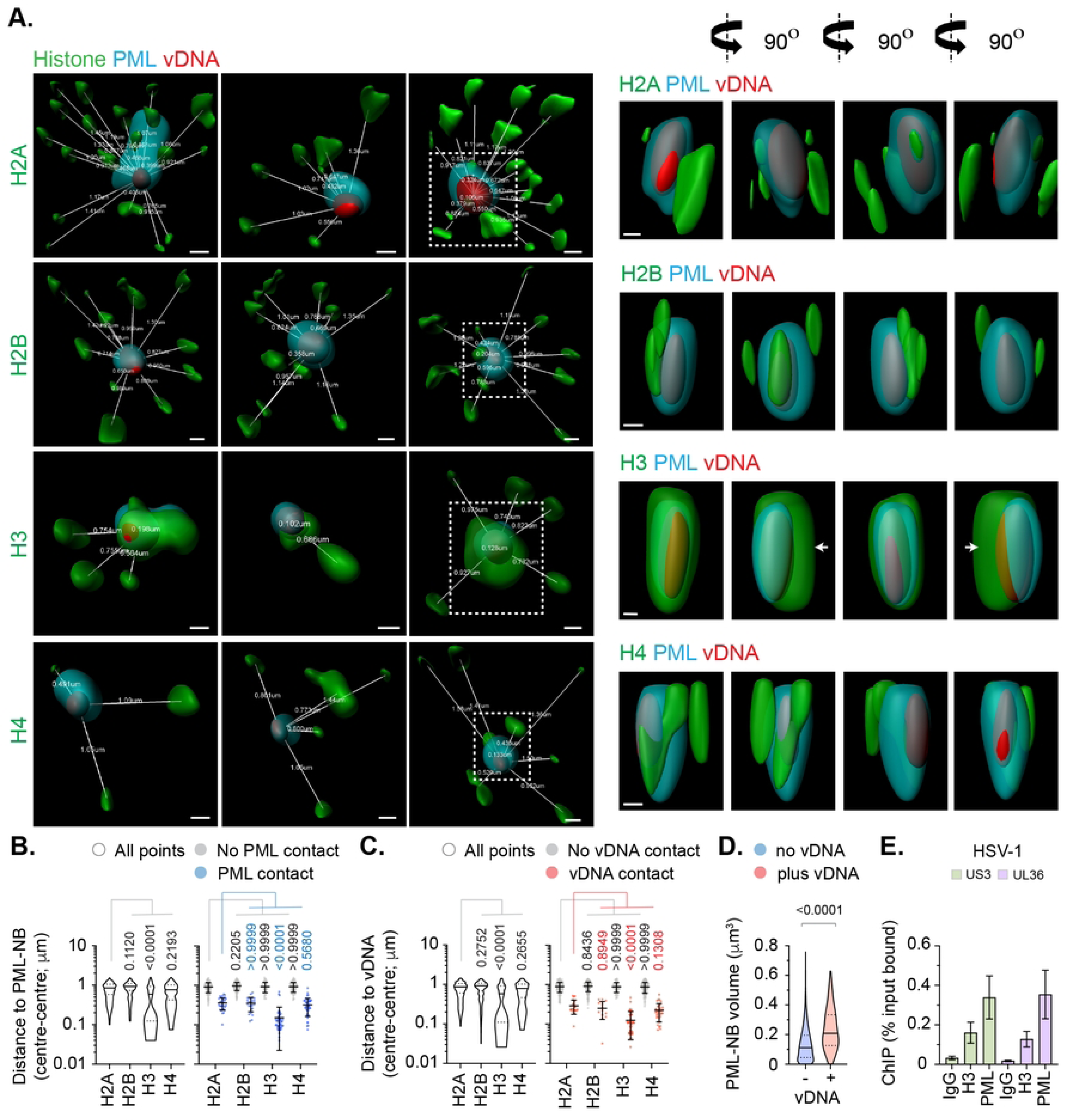
Cellular histones show alternate patterns of spatial proximity to nuclear infecting HSV-1 genomes. HFt cells were infected with WT HSV-1^EdC^ (MOI of 1 PFU/cell). Samples were fixed at 90 mpi and stained for proteins of interest (histones H2A, H2B, H3, or H4, and PML) by indirect immunofluorescence and vDNA by click chemistry. (A) Left; 3D render projections showing the spatial proximity and distance (white lines, μm) of endogenous cellular histones (green, as indicated) and PML (cyan) within a 2 μm^3^ region centred on vDNA (red). Right; 360° rotation of region of interest (dashed white boxes, left). Scale bars = 0.2 μm. (B/C) Quantitation of histone proximity (centre-to-centre distance, μm) to PML-NBs (B) or vDNA (C). Violin plots: median, solid black line; 25^th^ to 75^th^ percentile range, dotted lines. Scatter plots: black line, mean; whisker, SD. N>100 vDNA foci per sample condition (all points). One-way ANOVA Kruskal-Wallis test, *P*-values shown. (D) Quantitation of PML-NB volume (μm^3^) in the presence or absence of vDNA. N≥150 PML-NBs per condition. Mann-Whitney *U*-test, *P*-value shown. (E) HFt or HFt SPOT.PML.I expressing cells were infected with HSV-1 (MOI of 3 PFU/cell). Chromatin extracts were prepared at 90 mpi and subjected to ChIP using anti-histone H3, SPOT-tag, or species-matched IgG (negative control). Bound vDNA was quantified by qPCR using probes to HSV-1 US3 or UL36. Values were normalized to input loading controls and presented as percentage (%) input bound. Means and SEM shown. (A to E) Data derived from a minimum of three independent experiments. Raw values presented in S1 data.

### PML-NBs influence the spatial localization of histone H3.3 at vDNA

As PML-NB entrapment may exclude the *de novo* deposition of histones at vDNA, we next examined the colocalization of histones (H2A, H2B, H3.1/H3.3, and H4) with vDNA in PML knockout (KO) cells. Relative to non-targeting control (NTC) cells, over 80% of PML gRNA expressing cells lacked detectable PML-NBs (Fig. 5A, B). Infection of PML KO cells led to a reduction in the frequency of Daxx, histone H3, and histone H4 colocalization at vDNA relative to NTC cells (Fig. 5C, D, S7). Consistent with a PML-NB independent mechanism of Daxx recruitment to vDNA [16, 37, 56], a significant proportion of genomes retained colocalization with Daxx and histone H3 in PML KO cells (Fig. 5C, D). These data corroborate our HaCaT analysis (Fig. 3), demonstrating histone H3.3 and H4 to be actively recruited to vDNA upon nuclear infection independently of their native localization at PML-NBs. In contrast, histones H2A and H2B colocalization at vDNA remained below coincident threshold levels (Fig. 5C, D), demonstrating PML-NBs not to sterically inhibit the stable enrichment of these histones to vDNA. High-resolution imaging identified an increase in the spatial distance between histone H3 and vDNA between infected NTC and PML KO cells (Fig. 5E, F), identifying a role for PML in the spatial localization and/or stable enrichment of histone H3.3 at vDNA. We conclude that PML entrapment of HSV-1 genomes not to sterically inhibit the deposition of histones H2A or H2B at vDNA upon nuclear infection.

**Fig 5.**
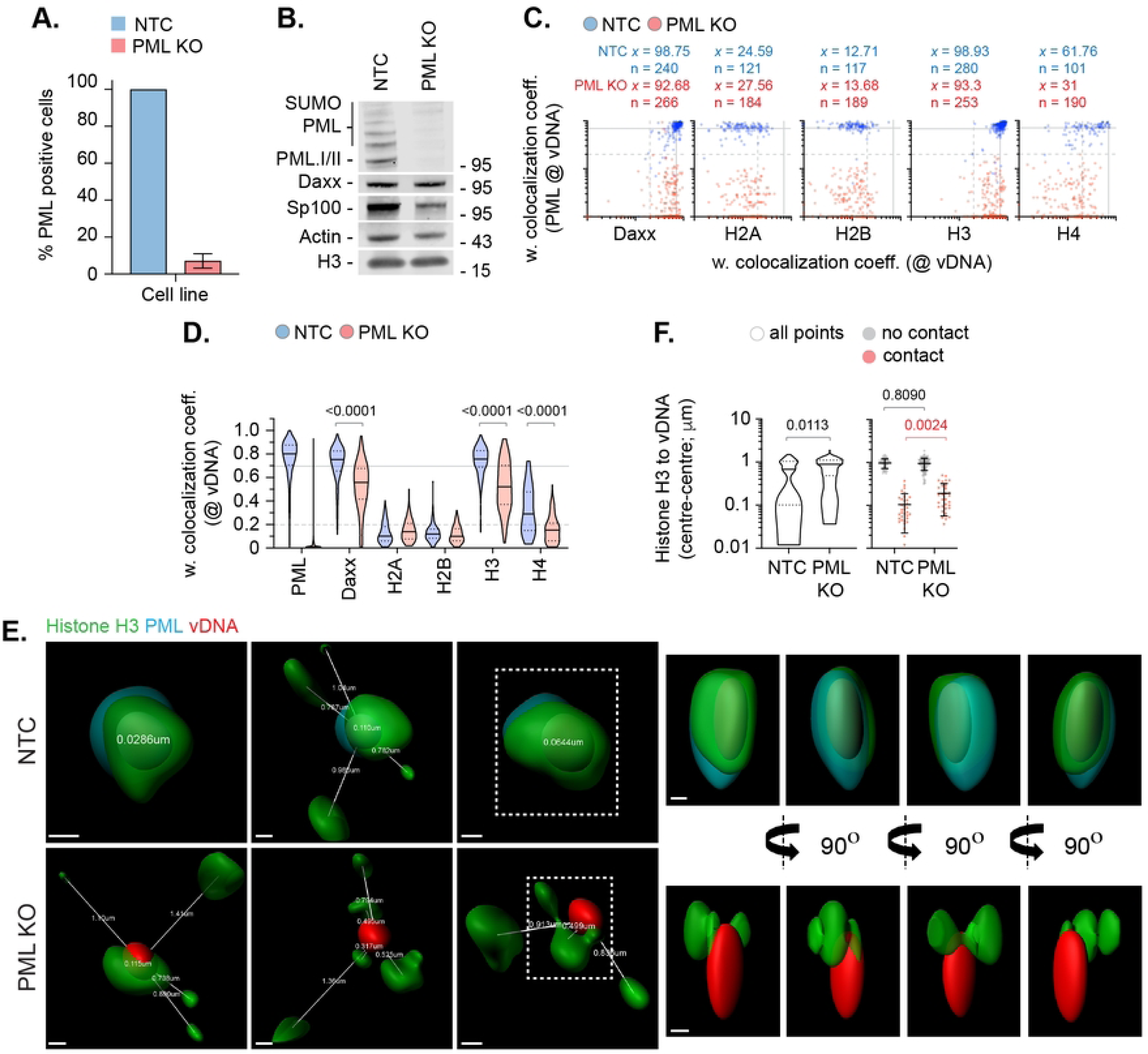
PML-NBs do not sterically inhibit histone H2A or H2B enrichment to nuclear infecting HSV-1 genomes. HFt cells were stably transduced with lentiviruses expressing CRISPR/CAS9 and non-targeting control (NTC) or PML-targeting (PML KO) gRNAs. (A) Quantitation of PML knockout in NTC and PML KO HFt cells by indirect immunofluorescence staining. N=17 fields of view per sample condition; means and SD shown. (B) Western blot of NTC or PML KO HFt WCLs. Membranes were probed for PML, Sp100, Daxx, Histone H3, and Actin (loading control). Molecular mass markers indicated. (C) NTC or PML KO HFt cells were infected with WT HSV-1^EdC^ (MOI of 1 PFU/cell). Cells were fixed at 90 mpi and stained for proteins of interest (Daxx, Histones H2A, H2B, H3, or H4, and PML) by indirect immunofluorescence and vDNA by click chemistry. Scatter plots showing paired weighted (w.) colocalization coefficient (coeff.) values of proteins of interest (indicated on *x*- and *y*-axis) at vDNA. Percentage (%) of genomes ≥ coincident threshold (*x/y* ≥ 0.2) per sample condition shown; number (n) of genome foci analysed per sample condition shown. Confocal images shown in Fig S7. (D) Distribution in protein colocalization frequency at vDNA (as in C). Violin plots: median, solid black line; 25^th^ to 75^th^ percentile range, dotted lines. Mann-Whitney *U*-test, *P*-values shown. (E) Left; 3D rendered projections of super-resolution images showing spatial proximity and distance (white lines, μm) of histone H3 (green) and PML (cyan) within a 2 μm^3^ region centred on vDNA (red). Right; 360° rotation of region of interest (dashed white boxes, left). Scale bars = 0.2 μm. (F) Quantitation of histone H3 proximity (centre-to-centre distance, μm). N≥90 vDNA foci per sample condition (all points). Violin plots (as described in D). Scatter plots: black lines, mean; whisker, SD. Mann-Whitney *U*-test, *P*-values shown. (A, C to F) Data derived from a minimum of three independent experiments. Raw values presented in S1 data.

### The histone H3.3/H4 chaperone Daxx restricts HSV-1 genome expansion

As Daxx is known to promote the deposition of histone H3.3/H4 at PML-NBs [49–51], we next examined the role of Daxx in the stable colocalization of Histone H3.3 at vDNA in NTC and Daxx KO cells. Relative to NTC cells, over 80% of Daxx gRNA expressing cells lacked detectable Daxx localization at PML-NBs (Fig. 6A-C, S8). Daxx KO led to a loss of ATRX, the binding partner of Daxx [53, 57], and histone H3.3 localization at PML-NBs (Fig. 6D, S8). Infection of Daxx KO cells led to a significant reduction in ATRX and histone H3.3 colocalization at vDNA relative to infected NTC cells (Fig. 6E, F, S9), demonstrating a Daxx-dependent mechanism of histone H3.3 enrichment at vDNA. High-resolution imaging identified PML-NB entrapment of vDNA to occur independently of Daxx (Fig. 6G) and for the spatial proximity of histone H3 at vDNA to increase between NTC and Daxx KO cells (Fig. 6H).

**Fig 6.**
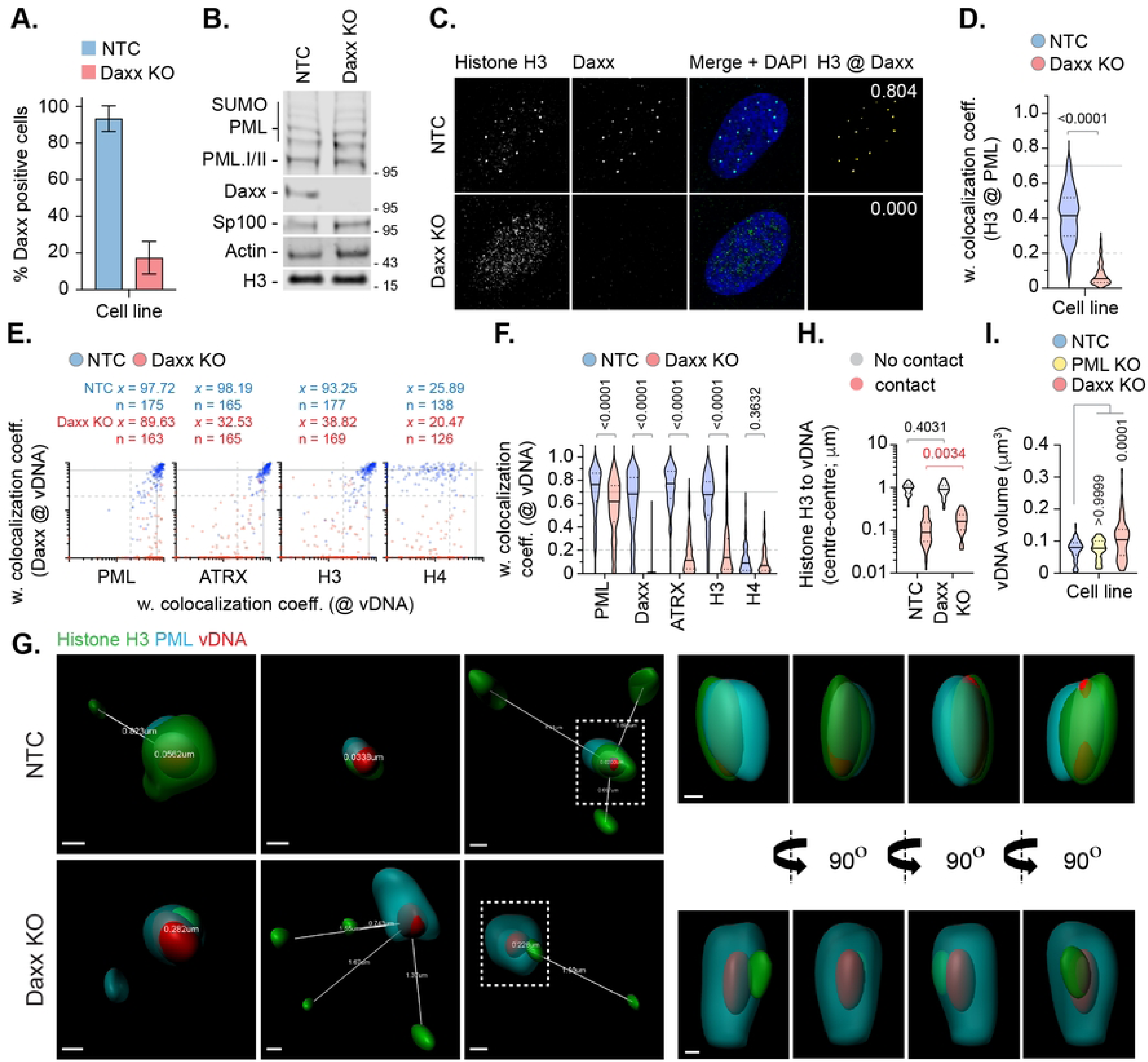
Daxx promotes histone H3 deposition at HSV-1 DNA to limit viral genome expansion. HFt cells were stably transduced with lentiviruses expressing CRISPR/CAS9 and non-targeting control (NTC) or Daxx-targeting (Daxx KO) gRNAs. (A) Quantitation of Daxx knockout in NTC or Daxx KO HFt cells by indirect immunofluorescence staining. N≥60 fields of view per sample condition; means and SD shown. Confocal images shown in Fig S8. (B) Western blot of NTC or Daxx KO HFt WCLs. Membranes were probed for PML, Sp100, Daxx, Histone H3, and Actin (loading control). Molecular mass markers indicated. (C) Confocal microscopy images of NTC or Daxx KO HFt cells stained for histone H3 (green) and Daxx (cyan). Nuclei were stained with DAPI (blue). Cut mask (yellow) highlights regions of colocalization between cellular proteins of interest; weighted (w.) colocalization coefficient (coeff.) shown. (D) Distribution in histone H3 colocalization frequency at PML-NBs in NTC and Daxx KO HFt cells. Violin plots: median, solid black line; 25^th^ to 75^th^ percentile range, dotted lines. N≥175 nuclei per sample condition. Mann-Whitney *U*-test, *P*-value shown. (E) NTC or Daxx KO HFt cells were infected with WT HSV-1^EdC^ (MOI of 1 PFU/cell). Cells were fixed at 90 mpi and stained for proteins of interest (PML, ATRX, histones H3 or H4, and Daxx) by indirect immunofluorescence and vDNA by click chemistry. Scatter plots showing paired w. colocalization coeff. values of proteins of interest (indicated on *x*- and *y*-axis) at vDNA. Percentage (%) of genomes ≥ coincident threshold (*x/y* ≥ 0.2) per sample condition shown; number (n) of genome foci analysed per sample condition shown. Confocal images shown in Fig. S9. (F) Distribution in protein colocalization frequency at vDNA (as in E). Violin plots (as described in D); Mann-Whitney *U*-test, *P*-values shown. (G) Left; 3D rendered projections of super-resolution images showing spatial proximity and distance (white lines, μm) of histone H3 (green) and PML (cyan) within a 2 μm^3^ region centred on vDNA (red) within infected NTC or Daxx KO HFt cells. Right; 360° rotation of region of interest (dashed white boxes, left). Scale bars = 0.2 μm. (H) Quantitation of histone H3 proximity (centre-to-centre distance, μm) to vDNA. Violin plots (as described in D). N≥80 vDNA foci per sample condition (all points); Mann-Whitney *U*-test, *P*-values shown. (I) Quantitation of genome volume (μm^3^) in NTC, PML KO, or Daxx KO HFt cells. N≥45 vDNA foci per sample condition; One-way ANOVA Kruskal-Wallis test, *P*-values shown. (A, C to I) Data derived from a minimum of three independent experiments. Raw values presented in S1 data.

Importantly, volumetric measurements of vDNA foci identified an increase in genome volume between Daxx KO and NTC or PML KO cells (Fig. 6I), indicative of change in vDNA decompaction previously ascribed to the initiation of viral transcription [17]. These data identify a role for Daxx in the deposition of chromatin at vDNA that limits the rate of HSV-1 genome expansion independently of the stable enrichment of histones H2A or H2B (Figs. 1 to 5).

### HSV-1 DNA retains a significant degree of compaction upon capsid release

As we had observed significant levels of population heterogeneity in the stable enrichment of histones H2A or H2B at vDNA (Figs. 1 to 5), these data suggested HSV-1 genome compaction upon nuclear infection to occur independently of stable canonical nucleosome assembly on a genome population basis. The visualization of herpesvirus genome entry into the nucleus remains poorly defined. However, electron microscopy (EM) and atomic force microscopy (AFM) studies have shown HSV-1 DNA to transit through the nuclear pore complex (NPC) as an electron-dense rod-shaped mass [13, 14]. In contrast, *in vitro* EM and AFM experiments have shown HSV-1 genomes to be released from viral capsids as linear strands of dsDNA when incubated in the presence of trypsin [58], detergent [59], or application of extreme pressure [60]. We hypothesized the presence of spermine in the capsid particle [9] and/or solvent density might be sufficient to retain genome compaction under less stringent conditions of release. To investigate, we performed *in vitro* genome release assays in the presence of non-specific carrier control protein (0.1 % FBS) to recapitulate the high protein content of the nucleoplasm (> 150 mg/ml; [61]) into which viral genomes are injected.

Genomes released under these conditions appeared as spherical foci (Fig. 7A; [16, 17]), as opposed to long filamentous strands of dsDNA. Volumetric measurements demonstrated these genomes to have equivalent dimensions to those observed inside infected cells at 90 mpi (median volume 0.08 *vs.* 0.0803 μm^3^, respectively; Fig. 7B). These data demonstrate HSV-1 genomes to retain a high degree of compaction post-capsid release independently of the presence of histones or chromatin modifying enzymes required to facilitate *de novo* nucleosome assembly. Taken together with our volumetric measurements in Daxx KO cells (Fig. 6I), these data support a role for Daxx to limit the rate of viral genome decompaction post-capsid release [14, 17], as opposed to promoting the re-compaction of linear DNA genomes into compact foci.

**Fig 7.**
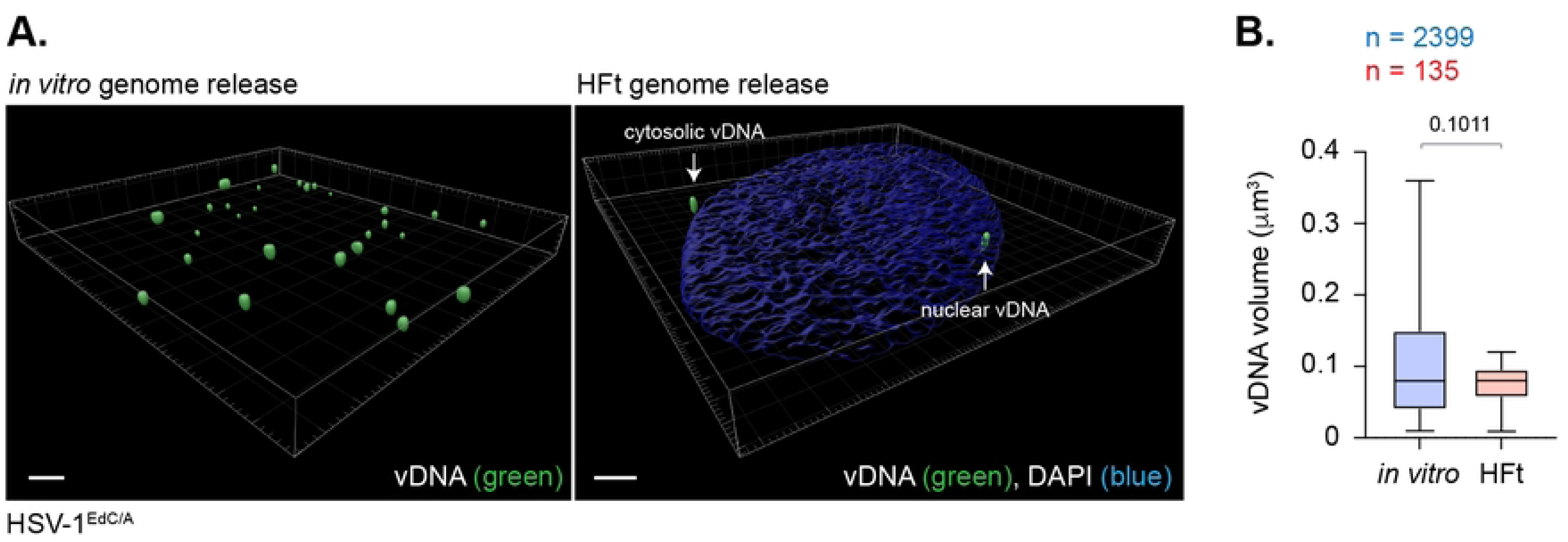
HSV-1 DNA retains a significant degree of native compaction post-capsid release. (A) 3D rendered projections of super-resolution images of HSV-1^EdC^ genomes released on glass coverslips *in vitro* or within infected HFt cells (MOI 1 PFU/cell) at 90 mpi (left and right-hand panels, respectively). vDNA was detected by click chemistry (green). Nuclei were stained with DAPI (blue). Scale bars = 2 μm. White arrows show cytosolic and nuclear vDNA foci. (B) Quantitation of vDNA foci dimensions (μm^3^) (as in A). Boxes, 25^th^ to 75^th^ percentile range; whisker, 5^th^ to 95^th^ percentile range; black line, median. Number (n) of genome foci analyzed per sample population shown; Mann-Whitney *U*-test, *P*-value shown. Data derived from a minimum of three independent experiments. Raw values presented in S1 data.

### The HSV-1 ubiquitin ligase ICP0 disperses Daxx and variant histone H3.3 from vDNA to stimulate genome expansion and the progression of IE transcription

We next examined the influence of ICP0 to disrupt the spatial localization of Daxx and variant histone H3.3 from vDNA entrapped within PML-NBs. HFt cells were infected with WT or ΔICP0 HSV-1^EdC/A^ (MOI of 0.5 PFU/cell) and fixed at 90 or 240 mpi. Consistent with the ICP0 dependent degradation of PML (Fig. S10A; [30, 35, 36]), PML colocalization at vDNA dropped below coincident threshold levels by 240 mpi during WT HSV-1 infection (Fig. 8A, B) [16]. An equivalent decrease in Daxx and variant histone H3.3 colocalization at vDNA could also be observed at this time point of infection, even though the stability of these proteins remained unchanged over the time course of infection (Fig. 8A, B, S10A). While the frequency of PML, Daxx, and histone H3.3 colocalization at vDNA decreased during ΔICP0 HSV-1 infection (Fig. 8A, B), a significant population of genomes retained colocalization of these proteins at vDNA (Fig. 8A; [16]). Volumetric measurements demonstrated WT HSV-1 genomes to undergo significant expansion by 240 mpi (Fig. 8C, D). In contrast, ΔICP0 HSV-1 genomes retained an equivalent degree of compaction to that observed at 90 mpi (Fig. 8C, D). Thus, the ICP0-dependent degradation of PML and dispersal of Daxx and variant histone H3.3 from vDNA stimulates the onset of viral genome expansion. Taken together with our Daxx KO analysis (Fig. 6), these data identify a role for Daxx in the formation of viral chromatin that limits the rate of viral genome decompaction upon nuclear infection.

**Fig 8.**
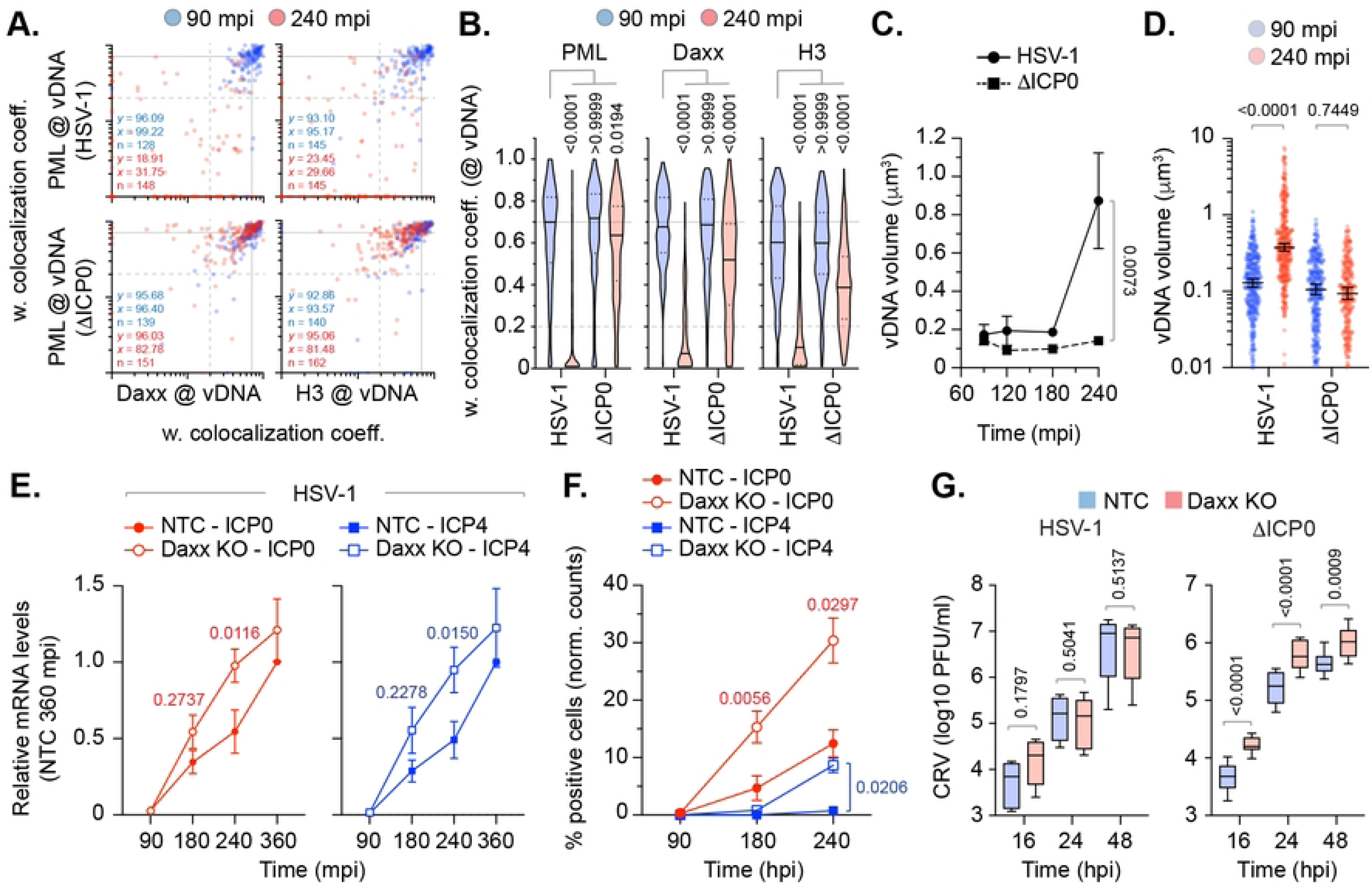
ICP0 disperses Daxx and variant histone H3.3 from vDNA to stimulate HSV-1 genome decompaction and progression of IE gene transcription. (A/B) HFt cells were infected with WT or ICP0 null-mutant (ΔICP0) HSV-1^EdC/A^ (MOI of 0.5 PFU/cell) for the indicated times (minutes post-infection; mpi). Cells were fixed and stained for Daxx, Histones H3, and PML by indirect immunofluorescence. vDNA was detected by click chemistry. (A) Scatter plots showing paired weighted (w.) colocalization coefficient (coeff.) values of proteins of interest (indicated on *x*- and *y*-axis) at vDNA at either 90 or 240 mpi (blue and red dots, respectively). Percentage (%) of genomes ≥ coincident threshold (*x/y* ≥ 0.2) per sample condition shown; number (n) of genome foci analysed per sample condition shown. (B) Distribution in protein colocalization frequency at vDNA (as in A). Violin plots: median w. colocalization coeff., solid black line; 25^th^ to 75^th^ percentile range, dotted black lines; coincidence threshold (0.2), dotted grey line; high confidence threshold (0.7), solid grey line. One-way ANOVA Kruskal-Wallis test, *P*-values shown. (C/D) Quantitation of HSV-1^EdC/A^ and ΔICP0^EdC/A^ DNA foci dimensions (μm^3^) over time (as indicated) in the presence of 2 μM supplemental EdC within the overlay medium; N≥270 genome foci per sample population. (C) Means and SD shown. Mann-Whitney *U*-test, *P*-value shown. (D) All points at 90 and 240 mpi shown. Mann-Whitney *U*-test, *P*-value shown.. (E) RT-qPCR analysis of HSV-1 IE gene transcription (ICP0 and ICP4, as indicated) in NTC and Daxx KO HFt cells infected with WT HSV-1 (MOI 0.5 PFU/cell) over time (mpi). Values were normalized to NTC infected cells at 360 mpi. Means and SEM shown; paired two-tailed t test, *P*-values shown. Individual replicate experiments shown in Fig. S10B. (F) Quantitation of HSV-1 immediate gene expression (ICP0 and ICP4) within AFP HSV-1 (MOI 0.5 PFU/cell) infected NTC and Daxx KO HFt cells over time (mpi) as determined by indirect immunofluorescence staining. Values normalized to the total population of DAPI positive cells (normalized [norm.] counts) and presented as percentage (%) positive cells. Means and SEM shown; paired two-tailed t test, *P*-values shown. (G) WT and ICP0 null-mutant (ΔICP0) HSV-1 cell supernatant titres derived from of infected NTC and Daxx KO HFt cells (MOI 0.5 PFU/cell) over time (hour post-infection; hpi). Boxes, 25^th^ to 75^th^ percentile range; whisker, 5^th^ to 95^th^ percentile range; black line, median. Mann-Whitney *U*-test, *P*-values shown. (A to F) Data derived from a minimum of three independent experiments. Raw values presented in S1 data.

To determine if this initial wave of chromatin assembly had a positive or negative impact on viral transcription, we next examined the kinetics of WT HSV-1 immediate-early (IE) transcription in NTC and Daxx KO cells. RT-qPCR analysis identified a delay in WT HSV-1 IE transcription (ICP0 and ICP4) between 90 and 240 mpi in NTC cells relative to Daxx KO cells, which began to recover to Daxx KO levels by 360 mpi (Fig. 8E, S10B). An equivalent delay in ICP0 and ICP4 protein expression was also observed in NTC cells relative to Daxx KO cells (Fig. 8F). These data demonstrate the Daxx mediated deposition of histones H3.3/H4 at vDNA to restrict the onset of WT HSV-1 IE transcription independently of the stable enrichment of histones H2A or H2B at vDNA (Figs. 1 to 5). Virus yield assays demonstrated no significant difference in WT HSV-1 titres between NTC and Daxx KO cells (Fig. 8G; HSV-1), consistent with a recovery in IE transcription by 360 mpi (Fig. 8E). In contrast, infection with ΔICP0 HSV-1 led to higher viral titres in Daxx KO relative to NTC cells (Fig. 8G; ΔICP0). Taken together, we conclude the histone chaperone properties of Daxx to play a key role in the assembly of viral chromatin that limits the rate of viral genome decompaction and progression of IE transcription. This host response to nuclear infection is antagonized by ICP0, which disperses Daxx from vDNA to destabilize this initial wave of replication-independent chromatin assembly to stimulate the progression of viral genome decompaction, IE transcription, and onset of HSV-1 lytic replication.

## Discussion

While the epigenetic modification of viral chromatin is known to play a fundamental role in the transcriptional regulation of herpesviruses during both lytic and latent phases of infection [23–25, 27], the prerequisite assembly of viral chromatin prior to its modification has remained poorly defined on a genome population basis. Here we utilise high-resolution quantitative imaging to investigate the population heterogeneity and spatial proximity of histones at vDNA upon HSV-1 nuclear infection at single-genome resolution.

Our imaging analysis identifies nuclear infecting HSV-1 genomes to asynchronously associate with canonical histones H2A, H2B, and H3.1 at a frequency significantly lower than that observed for PML-NB host factors (PML, Daxx, ATRX) or variant histone H3.3 at 90 mpi (Figs. 1 to 5). These data identify significant population heterogeneity in the recruitment of cellular histones to infecting HSV-1 genomes that is likely to differentially influence the kinetics or outcome of HSV-1 nuclear infection on an individual genome basis. Our data are consistent with ChIP and micrococcal nuclease studies that have reported only a small percentage of soluble input genomes to stably bind histones in a manner distinct from that of nucleosome arrays associated with cellular chromatin [29, 45–47, 62–65]. Thus, care needs to be taken when interpreting infection studies heavily reliant on ChIP, as the enrichment of viral genomes bound to specific histones or identified to carry specific epigenetic modifications may represent only a fraction of the total population of genomes under investigation. While microscopy studies come with experimental limitations too (e.g., the detection of low abundant antigens and epitope masking), we demonstrate equivalent trends in canonical histone localization to occur independently of cell type, levels of histone expression, or potential differences in antibody avidity on a genome population basis (Figs. 1 to 3). Importantly, our imaging analysis could readily identify changes in host factor recruitment and spatial proximity at vDNA under a variety of genetic (e.g., NTC *vs*. PML or Daxx KO; Figs. 5 and 6) and infection (WT *vs.* ΔICP0 HSV-1; Fig. 8) conditions, demonstrating the sensitivity of the approach employed. However, we cannot discount the possibility of highly transient or unstable histone interactions [66] that may be disrupted as a consequence of the microscopy conditions employed in our study. Our findings are broadly consistent with ATAC-Seq (assay for transposase-accessible chromatin sequencing) studies, which indicate that the majority HSV-1 DNA during lytic infection to lack protection from transposase activity by stable canonical nucleosomes [67, 68].

Our data support an alternate model of herpesvirus chromatin assembly and compaction, where vDNA retains a significant degree of compaction post-capsid release that progressively expands as infection progresses through viral genome decompaction (Fig. 6I, 7, 8C) [17]. Such a model could be explained by the presence of spermine within the capsid particle [9] and/or molecular crowding due to the density of the nucleoplasm into which vDNA is injected. We demonstrate HSV-1 genomes released *in vitro* to have equivalent volumetric dimensions to that observed within infected cells at 90 mpi independently of the presence of histones or chromatin modifying enzymes required to facilitate *de novo* vDNA compaction (Fig. 7). These findings are supported by AFM studies that have shown infecting HSV-1 genomes transiting through the NPC to appear as rod-like densities and mass spectrometry experiments that have shown isolated vDNA not to stably bind histones upon nuclear infection [14, 29]. Together with our imaging analysis, these data provide compelling evidence demonstrating HSV-1 genomes are delivered into the nucleus in a pre-existing semi-compact state independently of chromatin assembly. Thus, the pressure-driven release of herpesvirus genomes from the capsid particle into the nuclei of cells is unlikely to occur as a linear strand of dsDNA, as often depicted in the literature [23, 24]. Rather, genome release from the capsid is more likely to occur in a ‘globular-like’ fashion, as ejected vDNA meets the protein density and impedance of the surrounding nucleoplasm that is required to be displaced for genome exit to occur. Such observations are consistent with bacteriophage studies, which have shown exterior solvent density to promote vDNA compaction to facilitate full genome exit from the particle [69]. This model warrants further investigation, as genome decompaction is likely to represent a key stage in the infectious lifecycle of all herpesviruses that will be intimately linked to the initiation of viral transcription and gene accessibility (Fig. 8) [17].

While canonical histones (H2A, H2B, and H3.1) were not stably enriched at viral genomes, we consistently observed PML, Daxx, and variant histone H3.3 to be localized at vDNA on a genome population basis, indicative of an active host response to nuclear infection [16, 39]. Notably, a degree of population heterogeneity in the stable co-recruitment of these host factors to vDNA could also be observed (e.g., PML plus Daxx; Fig. 5). Thus, kinetic differences in the spatiotemporal assembly of viral and cellular protein complexes on vDNA will likely contribute to the probability of any individual genome successfully initiating a productive infection (e.g., ΔICP0; Fig 8) [16, 70]. This observation is supported by single-cell RNA-Seq studies, which have identified individual cells to express distinct profiles of viral and host transcription [71, 72]. Thus, the application of quantitative imaging provides a powerful tool to investigate the spatiotemporal relationship between pro- and anti-viral host factors that actively compete for vDNA binding to regulate the outcome of infection [29, 73–75].

Although high-resolution imaging identified canonical histones H2A and H2B to make surface contact with vDNA on an individual genome basis (Fig. 4), these occurrences were rare in frequency and unique in their spatial arrangements relative to PML or variant histone H3.3 (Fig. 4 to 6). Notably, we observed the asymmetric localization of variant histone H3.3 at PML-NBs (Fig. 4A, white arrows), which likely reflects the transition of histone H3.3/H4 heterodimers deposited at PML-NBs by Daxx into cellular chromatin undergoing active remodelling [49–51, 76, 77]. These data suggest that the majority of variant histone H3.3 localized at PML-NBs is unlikely to be associated with viral chromatin directly (Fig. 4, 5), highlighting the importance of studying viral-host interactions in the context of the three-dimensional microenvironment in which they occur. We demonstrate PML to be an accessory component of viral chromatin prior to its degradation by ICP0 (Fig. 4E, 8, Fig S10A) [30], confirming ICP0 capable of targeting multiple anti-viral host factors bound to vDNA for proteasomal degradation to stimulate the progress of infection [75]. Additional study is warranted to determine how PML interacts with vDNA due to its importance as a host restriction factor to multiple pathogens [78, 79]. We show PML-NBs not to sterically inhibit the enrichment of canonical histones H2A or H2B at vDNA (Fig. 5). Thus, we find no evidence to support a hypothesis that PML-NBs act as a site for canonical nucleosome assembly upon nuclear infection. However, we do not discount the possibility that PML-NBs may contribute to the sequential assembly of non-canonical or intermediate form(s) of viral (hetero)chromatin, as we identify a role for Daxx in the deposition of histones at vDNA that limits the rate of viral genome decompaction and IE transcription (Fig. 6I, 8D). Nor do we discount a role for PML-NBs in the establishment or maintenance of nucleosomal heterochromatin during other phases of infection (e.g., HSV-1 latency). Under such conditions, cell-type specific host factors and/or alternate patterns of immune regulation may influence the assembly and/or maintenance of viral heterochromatin at PML-NBs in the absence of key transactivating proteins (e.g., VP16 or ICP0) [43, 80–82]. Thus, it would be of interest to determine the relative population of genomes that are in association with canonical or variant histones within latently infected neurones, as diversity in nucleosome composition, epigenetic modification, or degree of heterochromatin compaction may differentially influence the frequency of HSV-1 reactivation on an individual genome basis.

While it remains to be determined as to when and where the assembly of canonical histone tetramers reported to bind vDNA occurs during HSV-1 lytic infection [66, 83, 84], our data are consistent with a model of sequential histone loading onto vDNA that is mediated by individual histone chaperones [54, 85]. Consistent with previous results [38], we failed to observe the stable recruitment of HIRA at vDNA at 90 mpi (Fig. 1, S3), demonstrating the replication-independent deposition of variant histone H3.3 and histone H4 at vDNA to be largely dependent on Daxx. Such a model of sequential chromatin assembly is supported by iPOND studies, which have shown chromatin regulators to bind vDNA at distinct stages of infection [29, 74]. This initial wave of replication-independent chromatin assembly by Daxx appears to be relatively short lived, as the ICP0-dependent disruption of PML-NBs is sufficient to disperse Daxx and variant histone H3.3 from vDNA independently of their gross degradation (Fig. 8, S10A). Notably, ATRX has been reported to promote the stability of viral heterochromatin upon infection of fibroblasts [28]. Thus, the Daxx dependent recruitment of ATRX to vDNA may also contribute to the stabilization of intermediary forms of viral chromatin entrapped within PML-NBs. This hypothesis is consistent with ChIP studies that have shown ICP0 expression to reduce histone deposition on vDNA [45] and for Daxx/ATRX to work cooperatively with PML in the epigenetic silencing of ΔICP0 HSV-1 [28, 43, 86]. Our imaging analysis provides spatial context to these studies and identifies histone H3.3/H4 deposition at vDNA to occur prior to the stable enrichment of histones H2A or H2B upon nuclear infection. It remains to be determined if other histone variants (e.g., macroH2A, H2A.Z, or H2A.X) may participate in this initial wave of replication-independent chromatin assembly. However, we note the expression of histone H2B variants (e.g., H2B.1 and H2B.W) to be highly cell-type specific (testis, oocyte, and zygote; <u>HistoneDB 2.0</u>) [87]. Thus, the lack of detectible histone H2B enrichment at vDNA (Fig. 1 to 5) suggests viral chromatin entrapped within PML-NBs is likely to be unstable, possibly accounting for its relative ease of disruption by ICP0 following PML degradation [45]. Our data are consistent with a model of sequential histone loading, where histone H3/H4 heterodimers are loaded onto cellular DNA by Daxx as a tetrasome prior to histone H2A/H2B incorporation [18]. Whether this initial wave of replication-independent viral chromatin assembly is subject to further histone assembly and/or epigenetic modification in the absence of ICP0 remains an open question for future research.

While we observed variable levels of endogenous histone H4 localization at vDNA, the ectopic expression of histone H4-mEm led to equivalent levels of enrichment to that observed for histone H3.3-mEm at vDNA (Fig. 2). Notably, Daxx is known to bind histone H3.3/H4 heterodimers in a conformation specific manner that restricts histone accessibility to other chaperones [54]. We posit the non-stoichiometric levels of endogenous histone H4 enrichment observed at vDNA likely relate to epitope masking for this histone-antibody combination. Such findings highlight the importance of utilising multiple confirmatory controls when interpreting microscopy data (Fig. 2, S5, S6). We demonstrate Daxx to mediate the enrichment of histone H3.3 at vDNA that correlates with a delay in viral genome decompaction and a restriction in IE transcription independently of the stable enrichment of histones H2A and H2B on a genome population basis. Due to the importance of Daxx in the cellular restriction of multiple pathogens, additional investigation is warranted to determine if Daxx influences the viral genome decompaction state of other viruses known to actively target Daxx to stimulate the progress of infection [88].

In summary, we demonstrate HSV-1 genome compaction upon delivery to the nucleus to occur independently of the stable enrichment of canonical histones H2A and H2B at vDNA on a genome population basis. We identify a role for the histone H3.3/H4 chaperone Daxx in the replication-independent assembly of viral chromatin at vDNA that restricts the rate of HSV-1 genome decompaction to limit the progression of WT HSV-1 IE transcription. This initial wave of chromatin assembly is disrupted by ICP0, which induces the degradation of PML to disperse Daxx and variant histone H3.3 from vDNA that stimulates the expansion of viral genomes, progression of IE transcription, and efficient initiation of HSV-1 lytic replication. Thus, we identify HSV-1 genome decompaction upon nuclear entry to play a key role in the transcriptional regulation and functional outcome of HSV-1 infection. Findings that are likely to be highly pertinent to the transcriptional regulation of many nuclear replicating herpesvirus pathogens.

## Materials and Methods

### Cells and drugs

Human osteosarcoma (U2OS; ECACC 92022711), human keratinocyte (HaCaT; AddexBio, T0020001), hTERT immortalized human Retinal Pigmented Epithelial (RPE-1; ATCC, CRL-4000), and hTERT immortalized human foreskin fibroblast (HFt; [89]) cells were grown in Dulbecco’s Modified Eagle Medium (DMEM; Life Technologies, 41966). Primary human embryonic lung fibroblasts (HEL 299; PHE, 87042207) were grown in Minimal Eagle Medium (MEM; Invitrogen, 21090-22) supplemented with 1% L-glutamine (Invitrogen, 25030024) and 1% sodium pyruvate (Invitrogen, 11360039). All media was supplemented with 100 units/ml penicillin, 100 µg/ml streptomycin (Life Technologies, 15140-122) and 10% foetal bovine serum (FBS; Life Technologies, 10270). TERT immortalisation was maintained in the presence of 5 µg/ml of Hygromycin (Invitrogen, 10687-010). Cells transduced with lentiviruses were maintained in media supplemented with 1 µg/ml Puromycin (Sigma-Aldrich, P8833) for selection or 0.5 µg/ml Puromycin for maintenance.

For the inducible expression of auto-fluorescent proteins, cells were treated with media containing 0.1 µg/ml doxycycline (Sigma; D9891). All cell lines were cultured and maintained at 37°C in 5% CO_2_.

### Generation of inducible mEmerald-tagged histone, SPOT-tagged PML, and NTC, PML, and Daxx knock-out (KO) cell lines

Plasmids expressing histones H2A, H2B, H3.1, H3.3, and H4 with C-terminal mEmerald (mEm) fluorescent tags were a gift from Michael Davidson and obtained from Addgene (Cat. # 54110, 56475, 54115, 54116, 54117, respectively). cDNAs were cloned into the doxycycline inducible lentiviral vector pLKO.TetO/R.eYFPnls [90] replacing the eYFPnls ORF. A plasmid expressing eYFP.PML.I was a gift from Roger Everett [91]. cDNA encoding PML.I was ligated in frame with oligos encoding SPOT-tag into the lentiviral vector pLKO.TetO/R.eYFPnls replacing the eYFPnls ORF. Inducible histone-mEm, eYFPnls, or SPOT.PML.I HFt cell lines were generated by lentiviral transduction as described [37]. Non-targeting control (NTC), PML KO, and Daxx KO HFt cell lines were generated as described [37, 92]. Target sequences: NTC, ATC GTT TCC GCT TAA CGG CG; PML, CAC CGC GGG TGT GTC TGC ACC TA G; Daxx CAC CGT CTA TGT GGC AGA GAT CCG G. plentiCRIPSR v2 (a gift from Feng Zhang) was obtained from Addgene (Cat # 52961). pVSV-G and pCMVDR8.91 were a gift from Didier Trono. Cells were transfected using Lipofectamine LTX with PLUS reagent (Invitrogen; 15338100) as per the manufacturer’s instructions.

### Viruses

Wild-type (WT) HSV-1 17*syn*+ (HSV-1), its ICP0-null mutant derivative *dl*1403 (ΔICP0; [93]), and auto-fluorescent protein (AFP) variant that expresses ECFP-ICP4 and EYFP-ICP0 (AFP4/0; [94]) were propagated in RPE cells and titrated in U2OS cells, as previously described [70]. The labelling and purification of single or double labelled HSV-1 virions with 5-Ethynyl-2’-deoxycytidine (EdC; Sigma-Aldrich, T511307) and 7-Deaza-7-ethynyl-2’-deoxyadenosine (EdA; Jena Bioscience, CLK-099) was performed as described [16, 38].

Briefly, RPE cells were infected with WT or ICP0-null mutant HSV-1 at a MOI of 0.001 or 0.5 PFU/cell, respectively, and incubated at 33 °C. 24 h post-infection (hpi), infected cells were pulse labelled with EdC/A (final combined concentration of 1 µM) every 24 h until extensive cytopathic effect (CPE) was observed. Supernatant containing EdC/A labelled cell-released virus (CRV) was clarified by centrifugation (1500 rpm for 10 min), filtered through a 0.45 µm sterile filter, and purified through a NAP-25 Sephadex column (GE Healthcare; 17-0852-01). Purified virus was titrated on U2OS cells [70]. For virus yield assays, cells were infected at the indicated multiplicity of infection (MOI) for 1 h at 37 °C prior to overlay with media. Media containing infectious cell released virus (CRV) was harvested from wells at the indicated time points and titrated on U2OS cells as described [70].

### Virion genome release assay

1x10^8^ PFU of HSV-1^EdC/A^ labelled and column purified virus in DMEM containing 10% FBS was diluted 1 in 2 in TE (20mM Tris-HCL pH 7.5, 1 mM EDTA) and incubated at RT for 30 mins prior a 1 in 5 dilution in 100% MeOH (final concentration of 70 % MeOH and 0.1 % FBS). 50 μl of the virion solution was applied to poly-D-lysine (Sigma-Aldrich, P7405) treated coverslips and incubation at 60 °C in a pre-heated oven for 15 mins. Coverslips were fixed in 1.8 % formaldehyde in PBS for 10 mins, washed twice in PBS, and blocked in filter sterilised PBS containing 2 % FBS for 30 mins at RT prior to click chemistry.

### Antibodies

Mouse primary Abs: HSV-1 Major capsid protein VP5 (DM165; [95]), HSV-1 ICP0 (11060; [96]), HSV-1 ICP4 (58s; [97]), PML (Abcam, ab96051), HIRA (Millipore, 04-1488), Daxx (AbD Serotec, MCA2143) and Actin (Developmental Studies Hybridoma Bank, 224-236-1-s). Rabbit primary Abs: Histone 2A (Abcam, ab18255), Histone 2B (Abcam, ab1790), Histone H3 (Abcam, ab1791), Histone H4 (Abcam, ab10158), Daxx (Upstate, 07-471), Sp100 (GeneTex, GTX131569), ATRX (Santa Cruz, H300), Actin (Sigma-Aldrich, A5060), PML (Jena Biosciences, ABD-030; Bethyl Laboratories, A301-167A), and GFP (Abcam, ab290). Nanobody: H2A-H2B dimer (Chromotek; Atto488). Secondary antibodies used for detection: Alexa-488 or -647 donkey anti-mouse or -rabbit (Invitrogen; A21206, A21202, A31573, A31571), DyLight-680 or -800 goat anti-mouse or -rabbit (Thermo; 35568, SA5-35571, 35518, 35521), and goat anti-mouse HRP (Sigma-Aldrich, A4416).

### Chromatin immunoprecipitation (ChIP)

Cells were seeded into 60 mm dishes at a density of 1.5x10^6^ cells/dish. 24 h post-seeding, cells were either infected or induced to express proteins of interest by the addition of doxycycline (0.1 μg/ml) for 16 h prior to infection (ΔICP0 MOI of 3 PFU/cell). Media was aspirated from infected cell monolayers at 90 mpi, chromatin crosslinked using shortwave UV irradiation (235 nm, 9k x 0.1 mJ/cm^2^) using a UVP CL-1000 shortwave Ultraviolet Crosslinker. Chromatin was extracted using a Chromatin extraction kit (Abcam, ab117152) as per the manufacturer’s guidelines, sheared using a Branson cup horn sonicator (10 pulses at 25 % amplitude, 25 secs on and 15 secs off), and clarified by centrifugation (12,000 rpm for 10 mins) at 4 °C. 10 μg of soluble chromatin (input) was immunoprecipitated using a ChIP magnetic-one step kit (Abcam, ab156907) using 1 μg of polyclonal ChIP-grade antisera or species-matched negative control IgG immune sera o/n at 4 °C on a rotary wheel. Magnetic beads were washed, proteinase-K treated, and DNA extracted following the manufacture’s guidelines.

### Quantitative PCR (qPCR) and reverse transcriptase quantitative PCR (RT-qPCR)

HSV-1 genomes isolated by ChIP were quantified by DNA amplification using Taqman Fast Universal PCR Master mix (Applied Biosystems, 435042) using target and control-specific primer/probe mixes (Table 1) in sealed MicroAMP Fast Optical 96-well plates (Applied Biosystems, 4346907 and 4360954) on a 7500 Fast Real-Time PCR system (Applied Biosystems); 1x cycle of 90 °C for 20 secs, 40x cycles of 95 °C for 3 min and 60 °C for 30 secs. Cycle threshold (Ct) values were normalized to their respective input controls and expressed as a percentage of soluble input. Viral transcript expression was quantified by RT-qPCR. RNA was isolated from infected cells using an RNAeasy Plus Kit (Qiagen, 74134), according to manufacturer’s instructions. RT was performed using TaqMan Reverse Transcriptase Reagents Kit (Life Technologies, N8080234) with oligo(dT) primers. cDNA was quantified (as above) using ICP0 or ICP4 and GAPDH (Life Technologies; 4333764F) primer/probe mixes (Table 1). Ct values were normalized to GAPDH using the ΔΔCt method and expressed relative to the indicated control treatment.

**Table 1.**
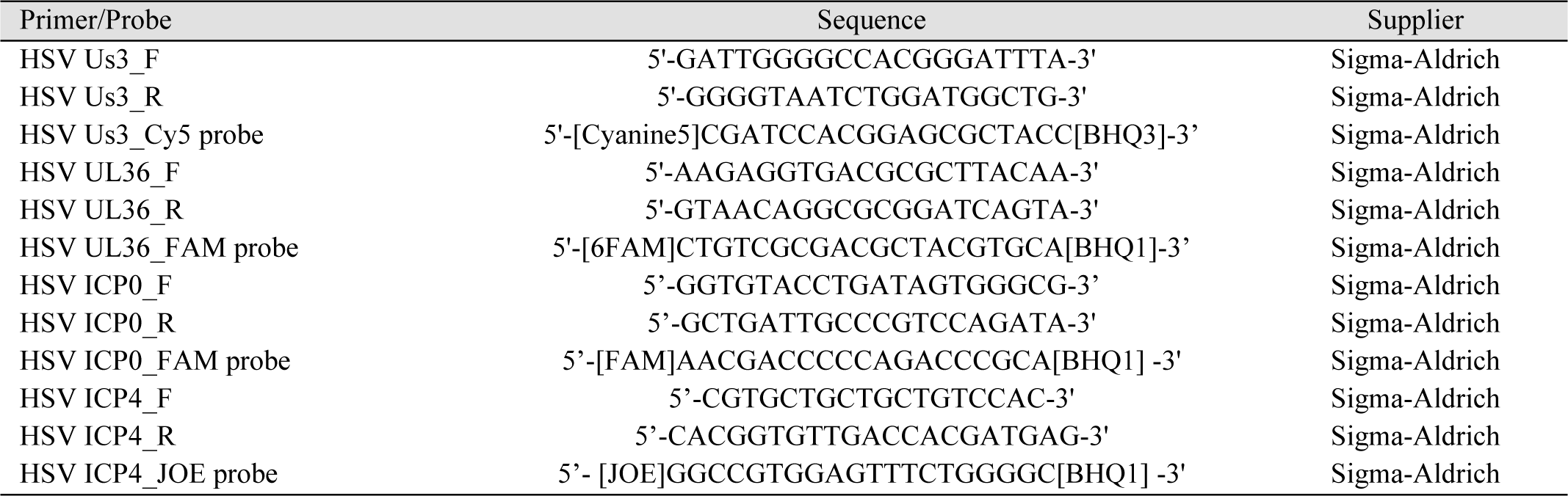
List of qPCR primer/probes used in the study.

### Western blot

Cells were washed twice in PBS before whole cell lysates were collected in 1x SDS-PAGE loading buffer supplemented with 2.5 M Urea (Sigma-aldrich; U0631) and 50 mM Dithiothreitol (DTT; Sigma-Aldrich, D0632). Proteins were resolved on 4-12% Bis-Tris NuPAGE gels (Invitrogen, NP0322BOX) in MES or MOPS buffer (Invitrogen, NP0001 or NP0002), and transferred to nitrocellulose (0.2 µm, Amersham; 15249794) using Novex transfer buffer (Invitrogen, NP0006-1) at 30 volts for 90 min. Membranes were blocked in PBS with 5% FBS for 1 h at RT. Membranes were incubated with primary antibodies diluted in blocking buffer at RT for at least 1 h or overnight at 4°C. Membranes were washed three times in PBST (PBS with 0.1 % Tween20), before incubation with secondary antibodies diluted in blocking buffer for 1 h at RT. Membranes were washed three times in PBST and rinsed in Milli-Q water before imaging on an Odyssey Infrared Imager (LiCor).

### Immunofluorescence and confocal microscopy

Cells were seeded onto 13 mm glass coverslips and incubated o/n at 37 °C in 5% CO_2_ prior to treatment or infection (as indicated). Cells were washed twice in CSK buffer (10 mM Herpes, 100 mM NaCl, 300 mM sucrose, 3mM MgCl_2_, 5 mM EDTA) prior to fixation and permeabilization in 1.8 % formaldehyde (Sigma-Aldrich, F8775) and 0.5 % Triton-X-100 (Sigma-Aldrich, T-9284) in CSK buffer for 10 mins at RT. Coverslips were washed twice in CSK buffer and blocked in 2 % Human Serum (HS; MP Biomedicals, 092931149) in PBS for 30 mins prior to click chemistry and immunostaining. Click chemistry was performed using a Click-iT-Plus EdU Alexa Fluor 488 or 555 imaging kit (ThermoFisher Scientific, C10637 or C10638) according to manufacturer’s instructions. For viral or host protein labelling, cells were incubated with primary antibodies diluted in PBS containing 2 % HS for 1 h, washed three times in PBS, and incubated in secondary antibodies and DAPI (Sigma-Aldrich, D9542) in PBS containing 2 % HS for 1 h. Coverslips were washed three times in PBS, twice in Milli-Q-H2O, and air dried prior to mounting onto Citiflour AF1 (Agar Scientific, R1320) on glass slides. Coverslips were examined using a Zeiss LSM 880 confocal microscope using the 63x Plan-Apochromat oil immersion lens (numerical aperture 1.4) using 405, 488, 545, and 633 nm laser lines. Zen black software was used for image capture, generating cut mask channels, and calculating weighted colocalization coefficients. High-resolution Z-series images were capture under LSM 880 Airy scan deconvolution settings using 1:1:1 capture conditions. Z-series images were process using Imaris (Bitplane v9.3) to produce rendered 3D images for distance and volumetric measurements.

### Statistical analysis

GraphPad Prism (version 10.2.2) was used for statistical analysis. For unpaired non-parametric data, a Kruskal-Wallis one-way ANOVA or Mann-Whitney *U*-test was applied. For unpaired parametric data, a two-tailed t test was applied. Statistical *P*-values are shown throughout. Significant differences were accepted at *P* ≤ 0.05.

## Acknowledgements

The authors would like to thank Mathew Weitzman for their constructive comments and input in the preparation of this manuscript.

## Funding

This work was funded by the Medical Research Council (MRC; https://mrc.ukri.org) grant MC_UU_12014/5 and MC_UU_00034/2 awarded to CB and the National Institute of Health (NIH) R01 NS105630 awarded to ARC. The funders had no role in the study design, data collection and analysis, decision to publish, or preparation of the manuscript.

## Data Availability

The data sets generated in this study are available in Supplemental File S1 data.

## Author Contributions

Conceptualization: APER, ARC, KLC, CB.

Methodology: APER, AO, VI, KLC, ZY, CB.

Investigation: APER, AO, VI, SMF, MCR, LO, ZY, CB.

Formal analysis: APER, AO, VI, SMF, MCR, LO, CB.

Data curation: APER, VI, SMF, MCR, LO, ZY, CB.

Project administration: APER, CB.

Supervision: CL, KLC, ARC, CB.

Resources: AO, SMF, KLC, CB.

Validating: APER, ZY, LO, CB

Funding acquisition: CB, ARC

Writing - original draft: APER, KLC, CB.

Writing - review editing: APER, MCR, IE, ARC, KLC, CB.

## Conflicts of Interest

The authors declare no conflict of interest.

## Supporting information

**Fig S1. Histone localization in mock-treated HFt cells.** Confocal microscopy images of data presented in Fig 1B. Mock-treated HFt cells were stained for Daxx, HIRA, histones H2A, H2B, H3, or H4 (Channel 1 [Ch.1]; green, as indicated) and PML (red) by indirect immunofluorescence. Nuclei were stained with DAPI (blue). Cut mask (yellow) highlights regions of colocalization between cellular proteins of interest and PML; weighted colocalization coefficient shown.

**Fig S2. Ectopic expression of fluorescently tagged histones in mock-treated HFt cells.** HFt cells were stably transduced with lentiviral vectors encoding C-terminally tagged fluorescent (mEmerald; mEm) histones or eYFPnls (negative control) as indicated. (A) Cells were induced to express proteins of interest for 24 h with doxycycline (DOX) prior to whole cell lysate (WCL) collection and western blotting. Membranes were probed for GFP and endogenous (endog.) histones H2A or H3. (B) RPE cells were transfected with plasmids expressing eGFP, H3.1-mEm, or H3.3-mEm for 24 h prior to WCL collection and western blotting. Membranes were probed for GFP and histone H3. (A/B) Molecular mass markers shown. (C to E) HFt cells were DOX induced for 6 h prior to fixation and indirect immunofluorescence staining for PML (red). Nuclei were stained with DAPI (blue). (C) Confocal microscopy images of histone-mEm or eYFPnls localization at PML-NBs. Cut mask (yellow) highlights regions of colocalization between cellular proteins of interest and PML; weighted (w.) colocalization coefficient (coeff.) shown. (D) Quantitation of the percentage of cells that demonstrate histone-mEm or eYFPnls colocalization at PML-NBs. Means and SD shown. (E) Violin plots showing histone-mEm w. colocalization coeff. frequency at PML-NBs: median w. colocalization coeff., solid black line; 25^th^ to 75^th^ percentile range, dotted black lines; coincidence threshold (0.2), dotted grey line; high confidence threshold (0.7), solid grey line. Mann-Whitney *U*-test, *P*-value shown. (D/E) N≥ 150 nuclei per sample condition. (A to E) Data derived from a minimum of three independent experiments. Raw values presented in S1 data.

**Fig S3. Localization of fluorescently tagged histones to mitotic cellular chromatin.** HFt cells stably transduced with lentiviral vectors encoding C-terminally tagged fluorescent (mEmerald; mEm) histones (as indicated) or eYFPnls (negative control) were induced with doxycycline for 6 h prior to fixation. Nuclei were stained with DAPI (blue). Representative x63 objective lens wide-field confocal microscopy images showing histone-mEm localization in mock-treated HFt cells. Dashed boxes show magnified regions of interest highlighting histone-mEm or eYFPnls localization at mitotic chromatin.

**Fig S4. Localization of endogenous histone to nuclear infecting HSV-1 genomes.** Confocal microscopy images of data presented in Fig. 1D to G. HFt cells were infected with WT HSV-1^EdC^ (MOI of 1 PFU/cell). Cells were fixed at 90 mpi and stained for Daxx, HIRA, histones H2A, H2B, H3, or H4 (Channel 1 [Ch.1]; green, as indicated) and PML (red) by indirect immunofluorescence. vDNA (red) was detected by click chemistry. Nuclei were stained with DAPI (blue). Cut mask (yellow) highlights regions of colocalization between cellular proteins of interest and vDNA or PML (as indicated); weighted colocalization coefficient shown. Dashed boxes show magnified regions of interest. White arrows highlight regions of colocalization at vDNA.

**Fig S5. Localization of endogenous histone H2A/H2B heterodimers to nuclear infecting HSV-1 genomes.** (A/B) Confocal microscopy images of data presented in Fig. 1E and F. HFt cells were mock-treated or infected with WT HSV-1^EdC^ (MOI of 1 PFU/cell). Cells were fixed at 90 mpi and stained for heterodimeric histone H2A/H2B (green) using a fluorescently conjugated nanobody and PML (cyan) by indirect immunofluorescence. vDNA (red) was detected by click chemistry. Nuclei were stained with DAPI (blue). Cut mask (yellow) highlights regions of colocalization between cellular proteins of interest and vDNA or cellular chromatin; weighted colocalization coefficient shown. Dashed box shows magnified region of interest. White arrows highlight regions of colocalization at vDNA. (B) Localization of histone H2A/H2B heterodimers to mitotic chromatin in mock-treated HFt cells.

**Fig S6. Localization of fluorescent histones to nuclear infecting HSV-1 genomes.** HFt cells stably transduced with lentiviral vectors encoding C-terminally tagged fluorescent (mEmerald; mEm) histones or eYFPnls (negative control) (Channel 1 [Ch.1]; green, as indicated) were induced with doxycycline for 6 h prior to infection with WT HSV-1^EdC^ (MOI of 1 PFU/cell). Cells were fixed at 90 mpi and stained for PML (cyan) by indirect immunofluorescence and vDNA (red) by click chemistry. Nuclei were stained with DAPI (blue). Cut mask (yellow) highlights regions of colocalization between cellular proteins of interest or vDNA (as indicated); weighted colocalization coefficient shown. Dashed boxes show magnified regions of interest. White arrows highlight regions of colocalization at vDNA.

**Fig S7. Localization of Daxx and endogenous histones to nuclear infecting HSV-1 genomes in NTC and PML KO HFt cells.** Confocal microscopy images of data presented in Fig. 5C, D. NTC and PML KO HFt cells were infected with WT HSV-1^EdC^ (MOI of 1 PFU/cell). Cells were fixed at 90 mpi and stained for Daxx, histones H2A, H2B, H3, or H4 (green, as indicated) and PML (cyan) by indirect immunofluorescence. vDNA (red) was detected by click chemistry. Nuclei were stained with DAPI (blue). Cut mask (yellow) highlights regions of colocalization between cellular proteins of interest and vDNA (as indicated); weighted colocalization coefficient shown. White arrows highlight regions of colocalization at vDNA. Dashed boxes show magnified regions of interest.

**Fig S8. Localization of histone H3, ATRX, and PML in NTC and Daxx KO HFt cells.** Confocal microscopy images of data presented in Fig. 6A. Mock-treated NTC and Daxx KO HFt cells were fixed and stained for PML, ATRX, and histone H3 (green, as indicated) and Daxx (cyan) by indirect immunofluorescence. Nuclei were stained with DAPI (blue). Cut mask (yellow) highlights regions of colocalization between cellular proteins of interest and Daxx (as indicated); weighted (w.) colocalization coefficient (coeff.) shown.

**Fig S9. Localization of histones H3 and H4 to nuclear infecting HSV-1 genomes in NTC and Daxx KO HFt cells.** Confocal microscopy images of data presented in Fig. 6E, F. NTC and Daxx KO HFt cells were infected with WT HSV-1^EdC^ (MOI of 1 PFU/cell). Cells were fixed at 90 mpi and stained for PML, ATRX, histones H3 or H4 (green, as indicated), and Daxx (cyan) by indirect immunofluorescence. vDNA (red) was detected by click chemistry. Nuclei were stained with DAPI (blue). Cut mask (yellow) highlights regions of colocalization between cellular proteins of interest and vDNA (as indicated); weighted colocalization coefficient shown. White arrows highlight regions of colocalization at vDNA. Dashed boxes show magnified regions of interest.

**Fig S10. Daxx restricts the progression of WT HSV-1 IE transcription.** (A) HFt cells were mock-treated or infected with WT or ICP0 null-mutant (ΔICP0) HSV-1 (MOI of 3 PFU/cell) in the absence or presence of the proteasome inhibitor MG132 (5 μM). WCLs were collected at the indicated times (h) post-infection (hpi) and analyzed by western blotting. Membranes were probed for ATRX, Daxx, PML, viral IE proteins (ICP0 and ICP4), histone H3, and actin (loading control). Molecular mass markers shown. < denotes the detection of a non-specific viral protein. (B) Independent replicate experiments of data presented in Fig. 8E. NTC and Daxx KO HFt cells were infected with WT HSV-1 (MOI 0.5 PFU/cell). RNA was extracted at the indicated times (minutes post-infection; mpi) and HSV-1 IE transcription (ICP0 and ICP4) quantified by RT-qPCR analysis. Values were normalized to infected NTC cells at 360 mpi. N=3 independent experiments. Means and SD per experiment shown. Raw values presented in S1 data.

**S1 Data.** Underlying data used for quantitative analysis in this study.

## Notes

### Competing Interest Statement

The authors have declared no competing interest.

